# Reprogramming of serine metabolism is an actionable vulnerability in *FLT3*-*ITD* driven acute myeloid leukaemia

**DOI:** 10.1101/2020.05.26.116392

**Authors:** Stefan Bjelosevic, Emily Gruber, Andrea Newbold, Lev M. Kats, Carolyn Shembrey, Thomas C. Abrehart, Izabela Todorovski, Giovanna Pomilio, Andrew H. Wei, Gareth P. Gregory, Stephin J. Vervoort, Kristin K. Brown, Ricky W. Johnstone

## Abstract

Activating FMS-like tyrosine kinase 3 (*FLT3*) mutations occur in approximately 30% of all acute myeloid leukaemias (AMLs) and are associated with poor prognosis. The limited clinical efficacy of FLT3 inhibitor monotherapy has highlighted the need for alternative therapeutic targets and treatments for FLT3-mutant AML. Using human and murine models of MLL-rearranged AML harbouring FLT3 internal tandem duplication (FLT3-ITD) and primary patient samples, we have demonstrated that FLT3-ITD promotes serine uptake and serine synthesis via transcriptional regulation of neutral amino acid transporters (*SLC1A4* and *SLC1A5*) and genes in the *de novo* serine synthesis pathway (*PHGDH* and *PSAT1*). Mechanistically, dysregulation of serine metabolism in FLT3-mutant AML is dependent on the mTORC1-ATF4 axis, that drives RNA-Pol II occupancy at *PHGDH, PSAT1, SLC1A4* and *SLC1A5*. Genetic or pharmacological inhibition of the *de novo* serine synthesis pathway selectively inhibited the proliferation of FLT3-ITD AML cells, and this was potentiated by withdrawal of exogenous serine. Purine supplementation effectively rescued the antiproliferative effect of inhibiting *de novo* serine synthesis, consistent with the idea that serine fuels purine nucleotide synthesis in FLT3-mutant AML. Pharmacological inhibition of the de novo serine synthesis pathway, using the PHGDH inhibitor WQ-2101, sensitises FLT3-mutant AML cells to the standard of care chemotherapy agent cytarabine via exacerbation of DNA damage. Collectively, these data reveal new insights as to how FLT3 mutations reprogram metabolism in AML, and reveal a combination therapy strategy to improve the treatment of FLT3-mutant AML.

**Statement of Significance:** FLT3 mutations are common in AML and are associated with poor prognosis. We show that FLT3-ITD stimulates serine metabolism, thereby rendering FLT3-ITD leukemias dependent on serine for proliferation and survival. This metabolic dependency can be exploited pharmacologically to sensitize FLT3-mutant AML to chemotherapy.

## INTRODUCTION

Understanding the genomic landscape of acute myeloid leukaemia (AML) has been facilitated through recent large-scale sequencing studies revealing the remarkable heterogeneity that underpins this aggressive disease (1-4). While new therapies for AML have recently been approved (5), the backbone of induction therapy in newly-diagnosed patients remains a high-dose chemotherapy regimen of the cytosine analogue cytarabine, in conjunction with an anthracycline such as daunorubicin. Long-term survival is relatively poor, with a 5-year survival rate of approximately 30% (6).

Activating mutations in receptor tyrosine kinases, such as the FMS-like tyrosine kinase 3 (*FLT3*) gene, are among the most frequently observed in AML, and are associated with dismal prognosis (7,8). FLT3 is primarily expressed on hematopoietic progenitor cells, and during early haematopoiesis coordinates a ligand-dependent signalling cascade that regulates the proliferation and maturation of the progenitor pool (9). FLT3 internal tandem duplication (ITD), the most common type of FLT3 mutation, promotes constitutive FLT3 tyrosine kinase activity and hyperactivation of downstream signalling pathways including JAK/STAT5, PI3K/mTOR and MAPK pathways (10-13). A number of FLT3 inhibitors have been developed and are in clinical trials. First generation inhibitors, such as midostaurin, (currently approved as frontline therapy in AML patients with FLT3-ITD mutations) sorafenib and lestaurtinib were developed as broad-spectrum kinase inhibitors. The inability of these agents to induce durable clinical responses resulted in the development of highly specific second-generation inhibitors (quizartinib, gilteritinib) with potent binding affinity for mutant FLT3. While clinical trials evaluating monotherapy of all the aforementioned agents have shown initial promise, the efficacy of FLT3 inhibitors is limited by rapid development of acquired resistance (14).

Extensive reprogramming of cellular metabolism is required to fulfil the increased energetic and biosynthetic demands associated with unrestrained proliferation and survival (15). Metabolic reprogramming also introduces new vulnerabilities that can be targeted for cancer therapy (16,17). In line with this notion, recent studies have demonstrated that FLT3-mutant AML cells are dependent on glycolysis and the pentose phosphate pathway (18,19). Moreover, glutamine metabolism has been shown to promote resistance to FLT3-targeted therapy (20,21). Emerging evidence suggests that many cancer cells are highly dependent on the non-essential amino acid serine (22,23). Serine is a major source of one-carbon units for the folate cycle, which provides metabolic intermediates for nucleotide synthesis and methylation reactions (24). The functional importance of serine metabolism in FLT3-mutant AML has not been defined.

Herein, we developed a physiologically relevant mouse model of MLL-rearranged, FLT3-mutant AML and used a multi-omics approach to identify actionable molecular vulnerabilities in this disease. We demonstrate that FLT3-ITD reprograms serine metabolism in a mTORC1-ATF4-dependent manner and show that mouse and human FLT3-mutant AML cells are selectively sensitive to genetic and pharmacological inhibition of serine metabolism. Importantly, this metabolic dependency can be exploited to sensitize FLT3-mutant AML cells to cytotoxic chemotherapy *in vitro* and *in vivo*. Collectively, these data reveal key insights into metabolic reprogramming events driven by FLT3 mutations in AML, and reveal a novel combinatorial therapeutic strategy to enhance the efficacy of standard of care chemotherapy in this aggressive subtype of AML.

## RESULTS

### A genetically engineered mouse model of *MLL*-rearranged, FLT3-ITD AML reveals FLT3-ITD is essential for leukaemia survival

Although FLT3-ITD mutations play a critical role in the pathogenesis of AML, alone they are insufficient to induce leukaemic transformation (25-27). We therefore generated a genetically engineered mouse model of doxycycline-inducible *FLT3-ITD* expression in *MLL-AF9* rearranged AML (28) (**Figure 1A**). Mice inoculated with haematopoietic progenitor cells co-expressing MLL-AF9 and inducible FLT3-ITD (denoted MLL-AF9/iFLT3-ITD) developed fully penetrant disease, detectable 7 days post transplantation via luciferase bioluminescence imaging (**Figure 1B, Figure 1C**). In contrast, detectable disease was only observed 35 days after mice were transplanted with MLL-AF9 expressing hemopoietic progenitor cells, although all transplanted mice did eventually succumb to disease (**Figure 1B, Figure 1C**). MLL-AF9/iFLT3-ITD cells expressing the dsRed reporter were harvested from bone marrow of mice at endpoint and used for subsequent experiments. We confirmed harvested cells faithfully recapitulated AML via immunophenotyping and histological analysis (**Supplementary Figure 1A**).

**Figure 1.**
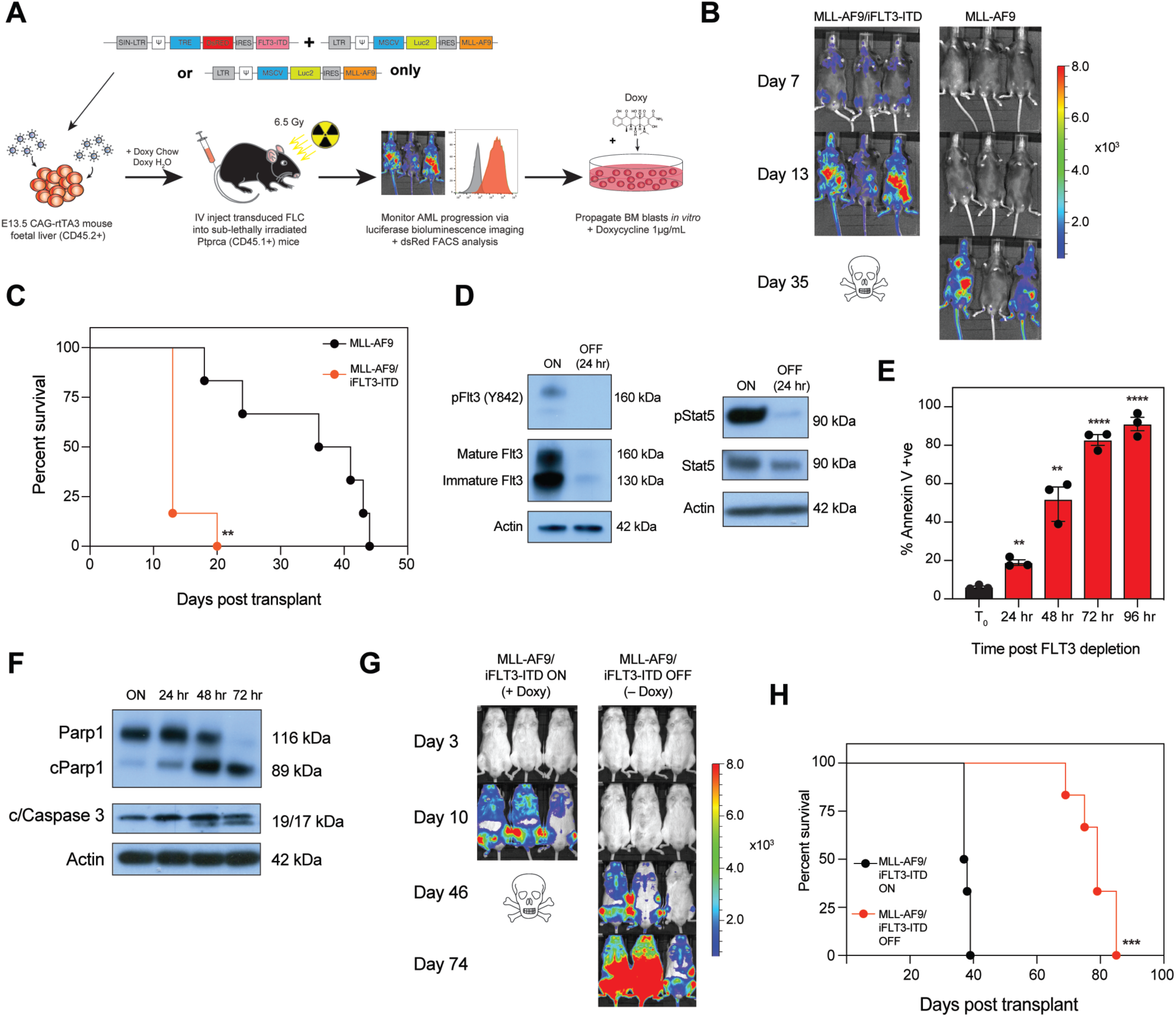
A genetically engineered mouse model of MLL-rearranged, FLT3-ITD AML reveals FLT3-ITD is essential for leukaemia survival. **A**. Schematic depicting generation of Tet-inducible FLT3-ITD mouse model. E13.5 murine foetal liver cells endogenously expressing the CAG-rtTA3 transgene were retrovirally transduced with Tet-DsRED-IRES-FLT3-ITD, MSCV-Luc2-IRES-MLL-AF9, or both vectors, and injected into sub-lethally irradiated *Ptprca* recipients. Leukaemia development was monitored via luciferase bioluminescence imaging and FACS analysis for dsRed, and at endpoint, bone marrow was harvested and blasts cultured *in vitro* in 1 μg/mL doxycycline. **B**. Primary transplant recipients were injected with 50 mg/kg D-luciferin and imaged in an IVIS bioluminescence imager to assess disease burden. **C**. Kaplan-Meier survival analysis of MLL-AF9/iFLT3-ITD or MLL-AF9 alone primary recipients. ** denotes *P* < 0.001 as determined by Mantel-Cox test, with *n =* 6 mice per condition. **D**. MLL-AF9/iFLT3-ITD cells were cultured *in vitro* with 1 μg/mL doxycycline and then washed twice in PBS with subsequent culture in doxycycline-free medium to deplete FLT3-ITD transgenic expression (MLL-AF9/iFLT3-ITD-OFF). Phosphorylated and total FLT3 and STAT5 expression was probed 24 hours post FLT3-ITD depletion via Western blot. β-Actin served as the loading control. **E**. FLT3-ITD was depleted in MLL-AF9/iFLT3-ITD cells as described above over a period of 96 hours, and percent apoptotic cells assessed every 24 hours via Annexin V staining and flow cytometry. ** denotes *P <* 0.01, \*\*\**P <* 0.001, **** *P <* 0.0001 by Student’s *t-*test. Error bars are representative of ± s.d. and three independent biological replicates. **F**. Apoptotic markers for cleaved/total PARP1 and cleaved/total Caspase 3 were assessed over 72 hours post FLT3-ITD depletion via Western blot in MLL-AF9/iFLT3-ITD cells. β-Actin served as the loading control. **G**. 1 × 10^6^ MLL-AF9/iFLT3-ITD cells were IV injected into secondary NSG recipients, and mice were placed on doxycycline-containing water and chow. After 72-hours, one arm of the study was placed on normal water and chow (MLL-AF9/iFLT3-ITD-OFF, – Doxy) and disease burden assessed via bioluminescence imaging. Mice were injected with 50 mg/kg D-luciferin and imaged as described above. **H**. Kaplan-Meier survival analysis of secondary recipients from **G**. *** denotes *P* < 0.001 determined by Mantel-Cox test, with *n =* 6 mice per condition.

To investigate the consequences of FLT3-ITD loss *in vitro*, MLL-AF9/iFLT3-ITD cells were propagated in doxycycline-free media for 24 hours (denoted MLL-AF9/iFLT3-ITD-OFF). Doxycycline withdrawal rapidly abolished FLT3-ITD expression, and resulted in decreased phosphorylation of the canonical FLT3 target STAT5 (**Figure 1D**). Importantly, FLT3-ITD depletion induced a time-dependent decline in cell viability, as determined by Annexin V staining (**Figure 1E, Supplementary Figure 1B**), immunoblot analysis of PARP/Caspase-3 cleavage (**Figure 1F**) and reduction in dsRed expression (**Supplementary Figure 1C**). To determine if loss of FLT3-ITD expression impairs *in vivo* disease progression, secondary NSG recipient mice were injected with MLL-AF9/iFLT3-ITD cells, and placed on doxycycline water and chow for 3 days prior to removal (**Supplementary Figure 1D**). Doxycycline withdrawal (and thus depletion of FLT3-ITD) dramatically reduced tumour burden (**Figure 1G**) and enhanced overall survival (**Figure 1H**). Depletion of FLT3-ITD following removal of doxycycline from leukemia-bearing mice was confirmed in bone marrow, spleen and peripheral blood using dsRed as a marker of transgene expression (**Supplementary Figure 1E**). Collectively these data demonstrate that MLL-AF9 and FLT3-ITD cooperate to drive onset and progression of AML and genetic depletion of FLT3-ITD alone is sufficient to induce apoptosis of MLL-AF9/iFLT3-ITD cells and extend overall survival in mice transplanted with these leukemias.

### *In vitro* and *in vivo* transcriptomics reveals *de novo* serine synthesis and serine uptake is regulated by FLT3-ITD

To elucidate the pathways involved in FLT3-ITD signalling in AML we performed 3’ RNA-sequencing comparing MLL-AF9/iFLT3-ITD cells cultured for 24 and 48 hours in the presence and absence of doxycycline (**Figure 2A**). We identified a total of 1,009 differentially expressed genes (DEGs) of which 575 were downregulated and 434 upregulated (*P* < 0.05 and –1 > logFC > 1) 24 hours post FLT3-ITD depletion (**Supplementary Table S1**). We further identified a total of 1,972 DEGs (1,057 downregulated, 915 upregulated, *P* < 0.05 and –1 > logFC > 1) 48 hours post FLT3-ITD depletion (**Supplementary Figure 2A; Supplementary Table S2**). Interestingly, among the most suppressed genes 24 hours after FLT3-ITD depletion were genes which encoded components of the *de novo* serine synthesis pathway (*Phgdh, Psat1*), neutral amino acid transporters involved in serine and glycine uptake (*Slc1a4, Slc1a5, Slc6a9*), the mitochondrial serine transporter (*Sfxn1)*, and one-carbon metabolism (*Mthfd2, Shmt1, Shmt2*) (**Figure 2B**). Indeed, terms that defined gene sets involved in serine biosynthesis, one-carbon metabolism and nucleotide metabolism were significantly suppressed following depletion of FLT3-ITD in MLL-AF9/iFLT3-ITD cells (**Figure 2C**). Examination of gene expression changes 48-hours post FLT3-ITD depletion generally showed exacerbation of loss of mRNA in all of the genes examined (**Supplementary Figure 2B**).

**Figure 2.**
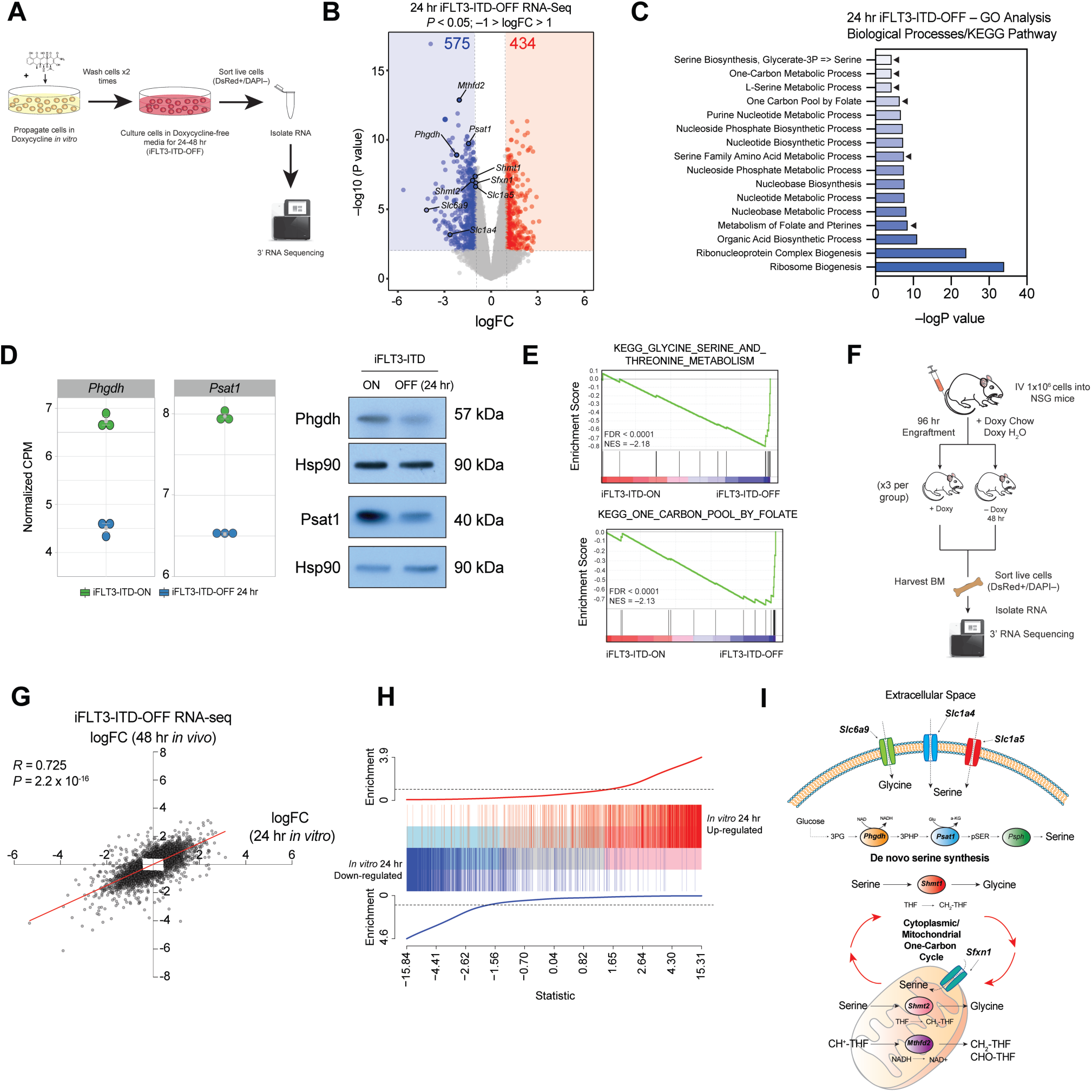
*In vitro* and *in vivo* transcriptomics reveals *de novo* serine synthesis and serine uptake is regulated by FLT3-ITD. **A**. RNA-sequencing of MLL-AF9/iFLT3-ITD cells post FLT3-ITD depletion. Cells were cultured *in vitro* in in 1 μg/mL doxycycline prior to 2 washes with PBS and subsequent culture for a further 24-48 hours in doxycycline-free medium. Cells were sorted for dsRed positive/DAPI negative cells, and RNA extracted. RNA was then 3’-RNA sequenced. **B**. Volcano plot of differentially expressed genes 24 hours post FLT3-ITD depletion, highlighting genes involved in *de novo* serine synthesis, serine and glycine uptake, and one-carbon/folate metabolism. Suppressed differentially expressed genes (575) are denoted by blue, and enriched genes (434) by red shading. Significance was defined as *P <* 0.05 and –1 > logFC > 1. **C**. KEGG pathway and biological processes GO term analysis of differentially expressed genes 24 hours post FLT3-ITD depletion. **D**. Dot plots of normalised counts-per-million (CPM) gene expression values 24-hours post FLT3-ITD depletion for *Phgdh* and *Psat1*, two of the most highly suppressed genes. Error bars denote ± s.d. and each condition was composed of 3 independent biological replicates. Protein expression of Phgdh and Psat1 were validated via Western blot 24 hours post FLT3-ITD depletion. Hsp90 served as the loading control. **E**. Enrichment score plots from GSEA using the KEGG signature for Glycine, Serine and Threonine Metabolism and One-Carbon Pool by Folate. FDR, false discovery rate; NES, normalised enrichment score. **F**. Schematic depicting *in vivo* experimental protocol. NSG mice were IV injected with 1 × 10^6^ MLL-AF9/iFLT3-ITD cells, and placed on doxycycline chow and water for 96 hours to allow for leukaemic engraftment. Mice were then placed on normal chow and water after 48 hours to allow for FLT3-ITD depletion (*n =* 3 mice per group). At 48 hours all mice were culled, blasts isolated from bone marrow, and sorted for dsRed positive/DAPI negative cells. RNA was isolated and 3’ RNA-sequenced. **G**. Correlation plot of differentially expressed genes post FLT3-ITD depletion at 24 hours (*in vitro*) and 48 hours (*in vivo*). Pearson’s correlation coefficient *R* = 0.725, *P <* 2.2 × 10^−16^. **H**. Barcode plot demonstrating enrichment of differentially expressed genes following 48 hour FLT3-ITD depletion *in vivo*, and the positive association with either significantly increased (red) (FDR < 0.05, logFC > 0.5) or significantly decreased (blue) (FDR < 0.05, logFC < 0.5) expressed genes post 24 hours post FLT3-ITD depletion *in vitro*. **I**. Schematic depicting biological pathways/functions of top-scoring genes involved in *de novo* serine synthesis, serine uptake, and one-carbon/folate metabolism.

Given the suppressed genes in our 24 hour *in vitro* transcriptomics data likely constitutes primary response genes, we performed all further analyses at this timepoint. Immunoblot analysis confirmed loss of Phgdh and Psat1 expression 24 hours following FLT3-ITD depletion, consistent with our RNA-seq data (**Figure 2D**). Gene set enrichment analysis (GSEA) further showed marked suppression of the KEGG pathway gene sets “glycine, serine and threonine metabolism”, and “one-carbon pool by folate”, consistent with GO Term analysis and individual gene changes observed (**Figure 2E**). It has been acknowledged that the use of tetracyclines (such as doxycycline) *in vitro* can have a direct effect on mitochondrial translation and cellular metabolism (29,30). To discount the possibility that differential gene expression was a confounding result of doxycycline withdrawal, we performed RNA-sequencing in our previously published Tet-inducible MLL-AF9 AML model (31). Analysis of RNA-seq data at 24 hours post MLL-AF9 depletion revealed minimal overlap with our FLT3-ITD depletion data. Importantly, the aforementioned genes were not differentially expressed in this system (**Supplementary Figure 2C; Supplementary Table S3**). These data provide important insights into the divergent molecular pathways regulated by FLT3-ITD and MLL-AF9. Our data indicates a key transcriptional role for FLT3-ITD in the regulation of important metabolic processes such as serine and one-carbon/folate metabolism, whilst MLL-AF9 transcriptionally regulates a well characterised network of genes involved in cellular differentiation (32). These observations are consistent with the biological consequences of FLT3-ITD and MLL-AF9 depletion, which results in induction of apoptosis as we have demonstrated and myeloid differentiation respectively (33).

To determine if our *in vitro* results were conserved *in vivo*, mice transplanted with MLL-AF9/iFLT3-ITD leukemias were divided into two cohorts, one with and one without doxycycline supplementation for 48 hours (**Figure 2F**). Leukemias were subsequently harvested from bone marrow for 3’-RNA-sequencing. Global transcriptome analysis showed a high degree of correlation between DEGs derived from 24 hour *in vitro* and 48-hour *in vivo* FLT3-ITD depletion (**Figure 2G; Supplementary Table S4**), with strong overlap in the direction and magnitude of DEGs observed (**Figure 2H**). Collectively, these data suggest that FLT3-ITD regulates *de novo* serine synthesis, serine uptake, and one-carbon metabolism (**Figure 2I**).

### Serine metabolism is a metabolic vulnerability in FLT3-ITD-driven AML

Serine is a nonessential amino acid that supports metabolic processes critical for the growth and survival of proliferating cells (23). To determine the functional role of serine metabolism in FLT3-ITD-driven human AML, we generated MV4-11 (*MLL-* rearranged, FLT3-ITD mutated) and OCI-AML3 (FLT3 wild type) AML cell lines in which *PHGDH*, the rate limiting step in the *de novo* serine synthesis pathway, was deleted using CRISPR-Cas9 gene editing (**Supplementary Figure 3A**). A competition assay in which *PHGDH*-deleted cells were co-cultured with wild-type cells and proliferation was monitored over 14 days was then performed. Deletion of *PHGDH* in MV4-11 cells, using two different sgRNAs, resulted in a loss of representation of these cells over time (**Figure 3A**). In contrast, deletion of *PHGDH* in OCI-AML3 cells had a minimal effect (**Figure 3A**). These data suggest that AML cells expressing mutant FLT3 may be dependent on *de novo* serine synthesis activity for growth and/or survival even when an exogenous supply of serine/glycine is available from the culture medium. To determine the role of exogenous serine in maintaining proliferation and/or survival of human leukaemia cells, MV4-11, OCI-AML3 and MOLM-13 (*MLL-*rearranged, FLT3-ITD mutated) cell lines were cultured in complete RPMI or serine/glycine-depleted RPMI for 72 hours. Proliferation across all cell lines was significantly impaired in serine/glycine-depleted conditions, suggesting exogenous serine is essential for optimal proliferation regardless of FLT3 mutational status (**Figure 3B**). Importantly, the viability of all cell lines was minimally affected following culture in serine/glycine-deficient media (**Figure 3B**).

**Figure 3.**
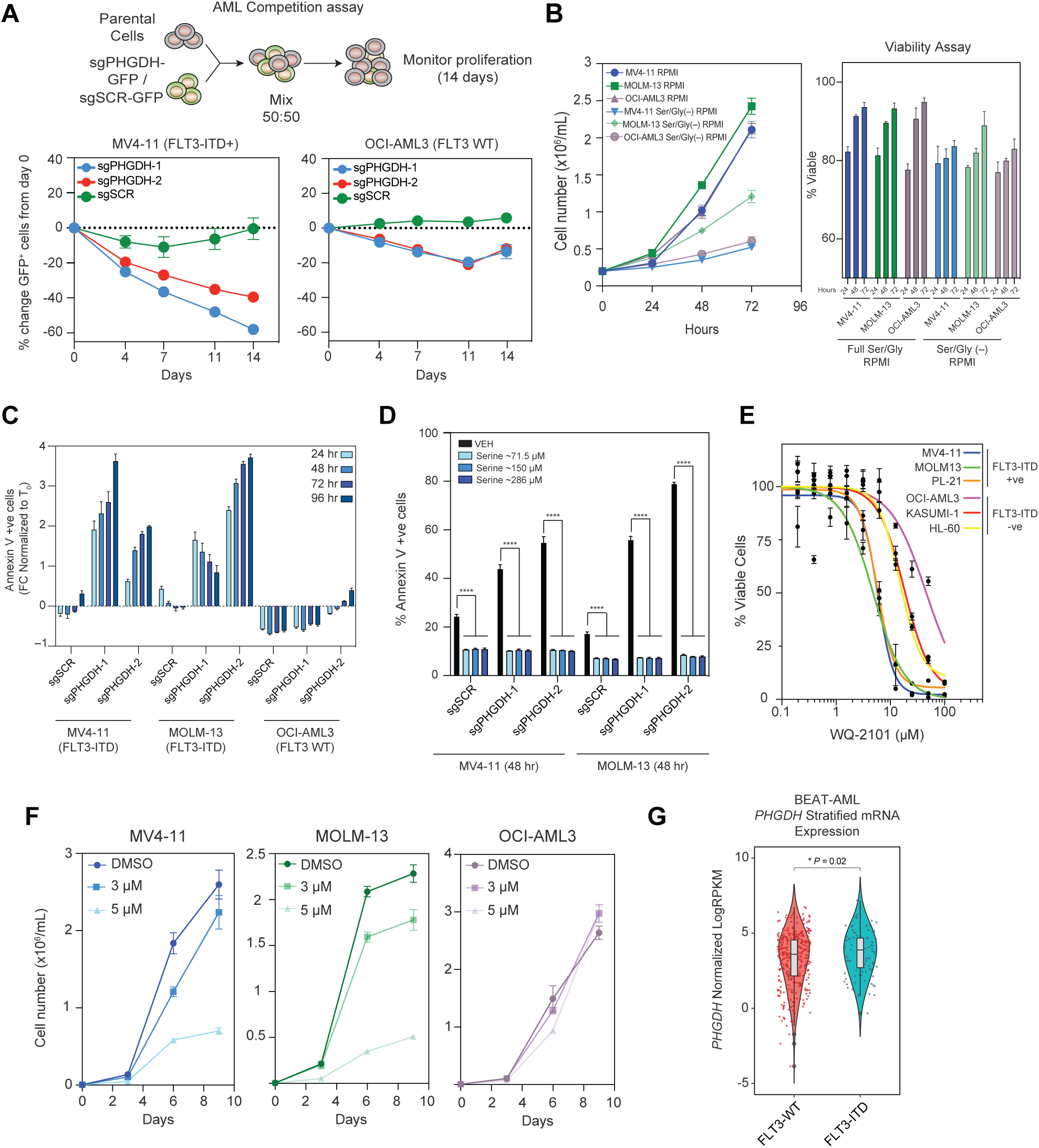
Serine metabolism is a metabolic vulnerability in FLT3-ITD-driven AML. **A**. Schematic depicting experimental protocol of a competition assay to determine if PHGDH depletion impairs cell proliferation. 50,000 GFP-positive cells transduced with either of two independent sgRNAs against PHGDH (sgPHGDH-1 or -2) or Scramble DNA (sgSCR) were co-cultured in a 1:1 ratio with parental non-transduced cells, and percentage change in GFP-positive cells relative to day 0 assessed at day 4, 7, 11 and 14. MV4-11 (FLT3-ITD mutant) or OCI-AML3 (FLT3 WT) cells were used, and cultured in complete RPMI. **B**. Cell counts assessing proliferation of two FLT3-ITD positive cell lines (MV4-11, MOLM-13) and FLT3 WT OCI-AML3 cells in either complete or serine/glycine-deprived RPMI over 72 hours. Cell viability was also determined via trypan-exclusion assays over the same period of time. **C**. MV4-11, MOLM-13 and OCI-AML3 transduced with either of two independent sgRNAs against PHGDH (sgPHGDH-1 or -2) or Scramble DNA (sgSCR) and were cultured in serine/glycine-deprived media for 96 hours, and cell viability determined via Annexin-V flow cytometry every 24 hours. Data presented as fold change relative to timepoint zero (T0). **D**. PHGDH-depleted MV4-11 and MOLM-13 cells were cultured in serine/glycine-deprived media for 48 hours. In each condition, three separate biologically relevant doses of serine (∼286 μM, serine concentration in commercial RPMI; ∼150 μM, serine concentration in peripheral blood; and ∼71.5 μM serine, representing a serine-insufficient state) were supplemented into culture medium at time of seeding. Cell viability was assessed at 48 hours via Annexin-V flow cytometry. **** *P <* 0.0001. Error bars are representative of ± s.d. and 3 independent biological replicates. **E**. Escalating doses of a small molecule inhibitor of PHGDH, WQ-2101, were screened against a panel of FLT3-ITD positive (MV4-11, MOLM-13 and PL-21) and FLT3 WT (OCI-AML3, Kasumi-1, HL-60), but otherwise genetically diverse, AML cell lines in complete RPMI. Viability was determined via Cell-Titer Glo. **F**. MV4-11, MOLM-13 and OCI-AML3 cells were treated with DMSO or two low doses of WQ-2101 (3 μM or 5 μM) and cell number assessed over a period of 9 days (timepoints at day 3, 6 and 9). All error bars are indicative of ± s.d., and 3 independent biological replicates. **G**. Violin plots illustrating normalised LogRPKM mRNA expression for *PHGDH* in FLT3 wild-type versus FLT3-ITD patients in the BEAT-AML dataset (*n* = 473). Statistical comparison is a Student’s *t-*test.

Utilising PHGDH-depleted MV4-11, OCI-AML3 and MOLM-13 cell lines (**Supplementary Figure 3A**), cell viability in serine/glycine-deficient media was then assessed. Strikingly, serine deprivation was selectively lethal to MV4-11 and MOLM-13 cell lines (**Figure 3C** and **Supplementary Figure 3B**), correlating with FLT3-ITD mutation status. Previous studies have shown that serine, but not glycine, supports one-carbon metabolism and proliferation of cancer cells (34). A rescue experiment was performed to determine if serine alone could reverse loss of viability observed following PHGDH depletion and serine/glycine withdrawal. Serine supplementation at three concentrations (∼286 μM, the concentration of serine in RPMI; ∼150 μM, the concentration of serine found in human serum (35); and 71.5 μM, to mimic a low serine state) was sufficient to reverse the apoptotic phenotype observed following global serine deprivation (**Figure 3D**), suggesting serine is essential for the survival of FLT3-ITD mutant AMLs. Collectively these experiments illustrate that, while exogenous serine is uniformly required for maximal proliferation of AML cells, depletion of *de novo* and exogenous serine is preferentially lethal to FLT3-ITD mutant AMLs.

To determine the effect of pharmacological inhibition of PHGDH in the context of AML, an allosteric small molecule inhibitor, WQ-2101 (36), was applied across a panel of FLT3-ITD mutant and FLT3 wildtype human AML cell lines. Potent induction of apoptosis was observed in cell lines harbouring FLT3-ITD mutations (MV4-11, MOLM-13, PL-21) at low micromolar ranges of WQ-2101, compared to FLT3 wildtype cell lines (OCI-AML3, Kasumi-1, HL-60) (**Figure 3E**). Moreover, a profound proliferative defect was observed in FLT3-ITD mutant cells exposed to sub-lethal doses of WQ-2101 (**Figure 3F** and **Supplementary Figure 3C**). Examination of the BEAT-AML dataset of 473 FLT3-ITD versus FLT3 wild-type patients (4) verified *PHGDH* is more highly expressed in FLT3-ITD-mutant patients, providing a potential mechanistic explanation for differential sensitivity to PHGDH inhibition (**Figure 3G**). Collectively, these data provide genetic and pharmacological evidence that disruption of the *de novo* serine synthesis pathway is preferentially toxic to FLT3-ITD-mutant AML.

### A FLT3-ITD/mTORC1/ATF4 axis transcriptionally modulates serine synthesis and serine transporters in FLT3-mutant AML

The identification of molecular pathways downstream of FLT3-ITD that regulate expression of genes involved in the *de novo* serine synthesis pathway, neutral amino acid transporters involved in serine/glycine uptake and one-carbon metabolism was then performed. To establish a transcriptional dataset of FLT3-ITD depletion in human cell lines, MV4-11 cells were exposed to the potent FLT3 inhibitor quizartinib (37,38) and gene expression profiling was performed by 3’-RNA sequencing (**Supplementary Table S5)**. Differentially repressed genes from quizartinib-treated human MV4-11 cells were compared to those detailed in Figure 2 from MLL-AF9/iFLT3-ITD cells cultured in the absence of doxycycline for 24 hours (**Figure 4A**). A total of 84 genes overlapped in both datasets and included *PHGDH, PSAT1, SLC1A4, SLC1A5, SLC6A9, SHMT2 and MTHFD2* (**Supplementary Table S6**). Consistent with the data in Figure 2D, showing that genetic depletion of FLT3-ITD in MLL-AF9/iFLT3-ITD cells resulted in reduced expression of Phgdh, a similar reduction in PHGDH protein expression was observed following quizartinib treatment of MV4-11 cells (**Supplementary Figure 4A**). To identify potential transcription factors mediating FLT3-ITD-dependent gene expression, the i-cisTarget motif analysis platform was used to interrogate the 84 overlapping genes (39). Strikingly, an ATF4 binding motif was identified as the most highly enriched (**Figure 4B**). ATF4 is a stress-induced master transcriptional regulator of amino acid metabolism and has previously been linked with regulation of serine metabolism (40). Indeed, ATF4 itself and several canonical ATF4 target genes (*ASNS, DDIT3* and *CHAC1*) were transcriptionally repressed following pharmacological inhibition of FLT3 in MV4-11 cells (**Figure 4C**), as well as in MLL-AF9/iFLT3-ITD cells 24 hours post FLT3-ITD depletion (**Supplementary Figure 4B**) with complete loss of ATF4 expression at the protein level (**Figure 4D**). Knockdown of ATF4 in MV4-11 cells resulted in a reduction in PHGDH and PSAT1 expression indicating ATF4 regulates the *de novo* serine synthesis pathway in FLT3-ITD mutant AML (**Figure 4E**).

**Figure 4.**
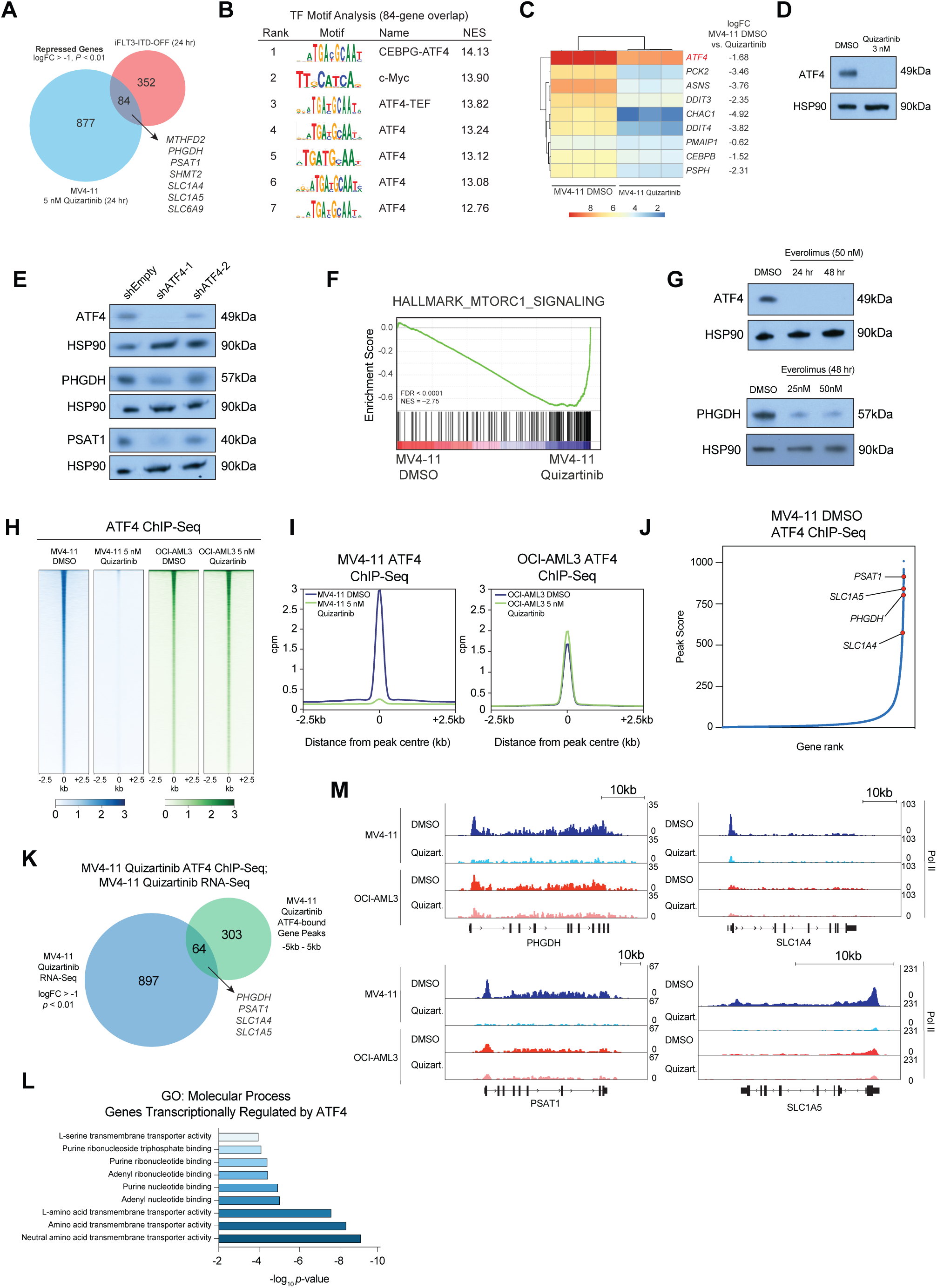
A FLT3-ITD/mTORC1/ATF4 axis transcriptionally modulates serine synthesis and serine transporters in FLT3-mutant AML. **A**. Venn diagram of suppressed DEGs from MLL-AF9/iFLT3-ITD-OFF cells at 24 hours and human MV4-11 suppressed DEGs in response to 5 nM quizartinib for 24 hours. A logFC > –1, *P* < 0.01 statistical cut-off was applied. **B**. Transcription factors with enriched binding sites in genes differentially suppressed upon FLT3-ITD depletion in genetic murine and human pharmacologic cell contexts (84-gene overlap from Figure 4A). Analysis is ranked using normalised enrichment score (NES) and was performed using the i-cisTarget motif analysis platform. **C**. Heatmap of ATF4 target gene suppression post treatment of MV4-11 cells with 5 nM quizartinib or DMSO for 24 hours. All gene expression changes are denoted as logFC. **D**. Western blot depicting rapid loss of ATF4 in MV4-11 cells post 24 hours of treatment with low-dose (3 nM) quizartinib. HSP90 acted as the loading control. **E**. Two independent shRNAs against ATF4 (denoted shATF4-1 and shATF4-2) and an empty vector control were lentivirally transduced into MV4-11 cells and induced in the presence of doxycycline for 72 hours prior to assessment of ATF4, PHGDH and PSAT1 protein expression via Western blot. HSP90 served as the loading control. **F**. Gene set enrichment (GSEA) plots depicting suppressed mTORC1 signalling after treatment of MV4-11 cells with 5 nM quizartinib for 24 hours, FDR < 0.0001, NES = −2.75. **G**. MV4-11 cells were treated with 50 nM of the mTORC1 inhibitor everolimus for 24 and 48 hours and ATF4 expression assessed via Western blot. Cells were subsequently treated with 25 nM or 50 nM of everolimus for 48 hours and PHGDH expression assessed via Western blot. HSP90 served as the loading control in both experiments. **H**. Occupancy heatmaps of genome-wide ATF4 peaks in the presence and absence of 5 nM quizartinib for 24 hours in MV4-11 and OCI-AML3 cells. **I**. Average profile of genome-wide ATF4 peaks in the presence and absence of 5 nM quizartinib in MV4-11 and OCI-AML3 cells. **J**. Plot ranking genes by peak score as determined by MACS2 and Homer software. Data depicts MV4-11 DMSO-treated cells, and genes relevant to *de novo* serine synthesis and serine/glycine uptake are highlighted. **K**. Venn diagram depicting ATF4 transcriptionally regulated genes. MV4-11 cells were treated with 5 nM quizartinib for 24 hours and suppressed genes from 3’ RNA-sequencing (logFC > –1, *P* < 0.01) were overlapped with ATF4-bound peaks (−5kb to 5kb) post quizartinib treatment. **L**. GO Term analysis of ATF4 transcriptionally regulated genes from the Venn diagram union of J. **M**. ChIP-seq reads for RNA Pol II mapped to the PHGDH, PSAT1, SLC1A4 and SLC1A5 loci in MV4-11 and OCI-AML3 cells treated with 5 nM quizartinib or DMSO for 24 hours.

Gene set enrichment analysis of the dataset from MV4-11 cells treated with quizartinib revealed mTORC1 signalling to be among the most suppressed gene sets (**Figure 4F**), a finding that was recapitulated in MLL-AF9/iFLT3-ITD cells 24 hours post FLT3-ITD depletion (**Supplementary Figure 4C**). The link between FLT3-ITD and mTORC1 has been previously studied (41) with mTORC1 regulating both the stability of ATF4 mRNA transcripts and ATF4 translation through three upstream open reading frames (uORFs) in the 5’ UTR of ATF4 (42). Consistent with these observations, genetic and pharmacological inhibition of FLT3-ITD in MLL-AF9/iFLT3-ITD and MV4-11 cells respectively resulted in dephosphorylation of Ribosomal Protein S6, a canonical mTORC1 target (**Supplementary Figure 4D)**. Moreover, treatment of MV4-11 cells with the mTORC1-specific inhibitor everolimus potently ablated ATF4 and PHGDH expression (**Figure 4G**).

To determine how ATF4 modulates *de novo* serine synthesis and serine uptake, chromatin immunoprecipitation sequencing (ChIP-seq) was performed in FLT3-ITD positive MV4-11 cells, and FLT3 wild-type OCI-AML3 cells using an anti-ATF4 antibody. Acute 24-hour treatment of MV4-11 cells with quizartinib resulted in global loss of ATF4 occupancy across the genome (**Figure 4H, Figure 4I**) whilst ATF4 occupancy was not globally altered in quizartinib-treated OCI-AML3 cells (**Figure 4H** and **Figure 4I**). Interestingly, a greater degree of ATF4 occupancy was seen in the MV4-11 cells at basal conditions, despite greater total ATF4 protein expression in OCI-AML3 cells (**Figure 4I** and **Supplementary Figure 4E**). Peak calling analysis of ATF4-ChIP-seq data and subsequent annotation of peaks to genes in DMSO-treated MV4-11 cells identified 11,869 unique genes (5kb either side of TSS) while in quizartinib-treated MV4-11 cells this was reduced to 367 genes, with a 332-gene overlap (**Supplementary Figure 4F**). The 332 genes that remained bound by ATF4 in the presence of quizartinib maintained enrichment for the ATF4 binding motif (**Supplementary Figure 4G**) and comprised key components of the *de novo* serine synthesis pathway, namely *PHGDH* and *PSAT1*, and the channel transporters *SLC1A4* and *SLC1A5*, suggesting these genes are particularly strongly bound by ATF4. Ranking of all genes according to their ATF4 occupancy revealed that globally ATF4 binding was enriched at *PHGDH, PSAT1, SLC1A4* and *SLC1A5*, supporting the notion that these genes are strongly bound by ATF4 (**Figure 4J**).

To determine target genes directly transcriptionally regulated by ATF4, the RNA-seq data from Figure 4A was integrated with the ChIP-seq data from Supplementary Figure 4G. This analysis yielded 64 genes that were most highly enriched for ATF4 binding and were significantly repressed following treatment of MV4-11 cells with quizartinib (**Figure 4K; Supplementary Table S7**). GO-term analysis revealed a recurrent amino acid transmembrane transporter activity signature that included the serine pathway as well as a purine nucleotide binding signature (**Figure 4L**), consistent with a recent report that mTORC1 enhances incorporation of one-carbon units into purine nucleotide biosynthesis (43).

To confirm that FLT3-ITD functionally promotes transcriptional activity at genes involved in serine synthesis and serine uptake, ChIP-sequencing using an anti-RNA polymerase II (Pol II) antibody was performed in MV4-11 and OCI-AML3 cells cultured in the presence or absence of quizartinib for 24 hours (**Figure 4M**). In all instances, MV4-11 cells demonstrated greater Pol II occupancy in transcribed regions of *PHGDH, PSAT1, SLC1A4* and *SLC1A5* genes than corresponding regions in OCI-AML3 cells. Strikingly, quizartinib treatment almost completely ablated Pol II occupancy at these regions in MV4-11 cells, suggesting FLT3-ITD inhibition effectively silences active transcription of these genes. Together with our previous evidence that ATF4 knockdown reduces PHGDH and PSAT1 expression (Figure 4E), these data suggest that FLT3-ITD regulates *de novo* serine synthesis and serine uptake activity in an ATF4-dependent manner.

### FLT3-ITD regulates global serine metabolism and drives purine nucleotide biosynthesis

To determine if total serine abundance is altered upon FLT3-ITD or PHGDH inhibition, steady-state metabolome profiling was performed. A dramatic reduction of intracellular serine and glycine abundance was observed following treatment of MV4-11 cells with quizartinib (**Figure 5A**). In contrast, inhibition of *de novo* serine synthesis with WQ-2101 only marginally reduced global serine/glycine abundance (**Figure 5A**). Critically, serine/glycine levels in cells exposed to WQ-2101 and deprived of exogenous serine/glycine largely phenocopied the results seen following treatment of MV4-11 cells with quizartinib, providing evidence that FLT3-ITD inhibition suppresses both *de novo* serine synthesis and serine uptake (**Figure 5A**). Importantly, assessment of the abundance of other amino acids (**Supplementary Figure 5A and 5B**) demonstrated that pharmacological inhibition of FLT3-ITD or PHGDH did not induce global changes in amino acid abundance. Quizartinib treatment resulted in profound changes in metabolites associated with glycolysis, namely increased glucose-6-phosphate and fructose-6-phosphate, coupled with decreased fructose-1,6-bisphosphate (**Supplementary Figure 5C**), suggesting a block in glycolytic metabolism. Whilst this phenomenon is not necessarily related to specific effects on serine metabolism, it is indicative of disruption of central carbon metabolism as previously described (18,19). Furthermore, FLT3-ITD inhibition with quizartinib or inhibition of PHGDH with WQ-2101 resulted in dramatic alterations in the abundance of TCA cycle intermediates (**Supplementary Figure 5D**). Of note, PHGDH inhibition significantly increased the abundance of citrate/isocitrate and succinate, consistent with previous reports (44). To functionally examine serine uptake dynamics in AML cells, MV4-11, MOLM-13 and OCI-AML3 cells were pre-treated with quizartinib or WQ-2101 for 24 hours, and the uptake of radiolabelled serine was measured. FLT3-ITD inhibition potently reduced serine uptake in FLT3-ITD positive MV4-11 and MOLM-13 cells (**Figure 5B**). In contrast, no change in serine uptake was observed in FLT3 wild-type OCI-AML3 (**Figure 5B**).

**Figure 5.**
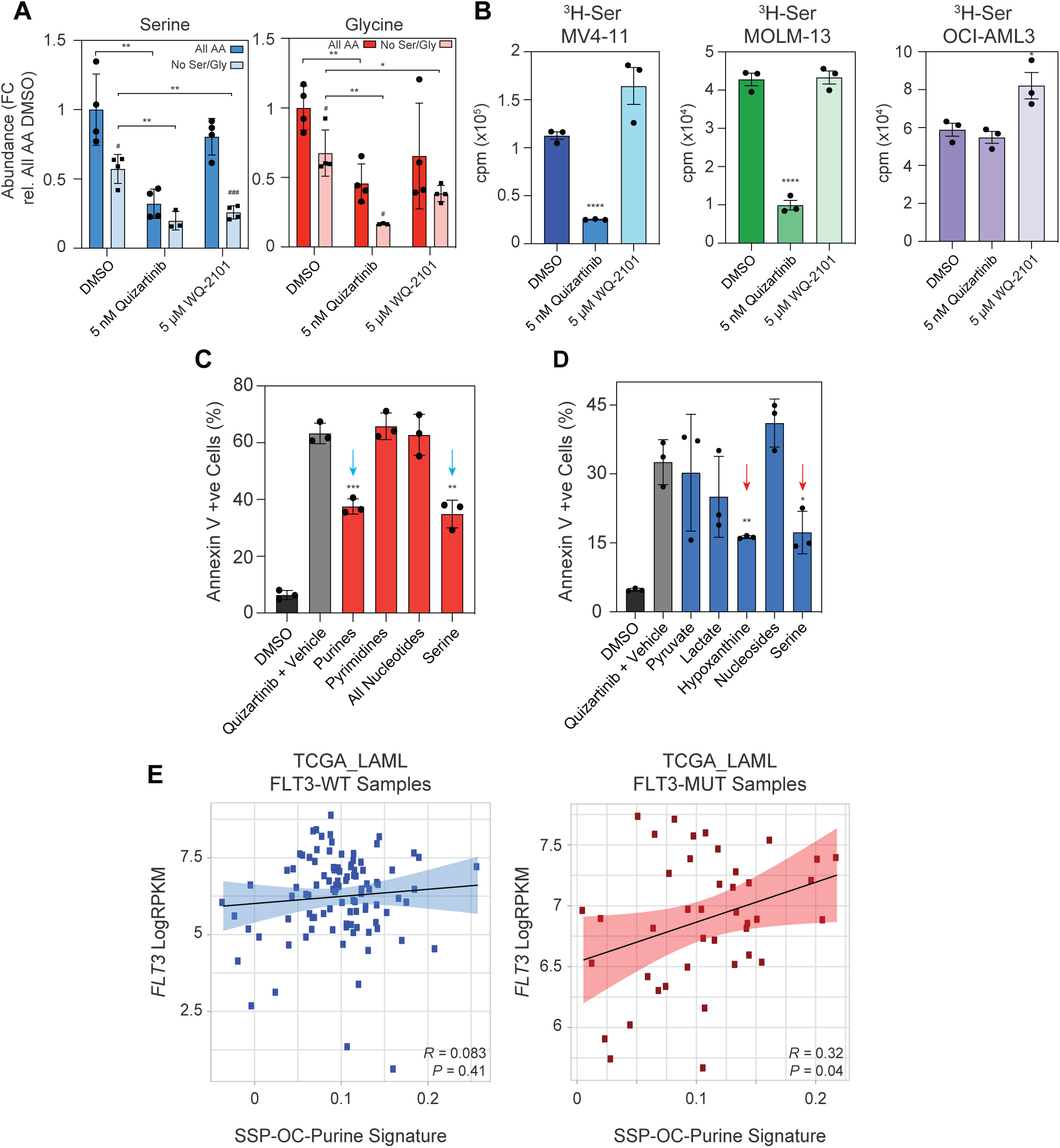
FLT3-ITD regulates global serine metabolism and drives purine nucleotide biosynthesis. **A**. Fold changes in serine and glycine abundance, as measured by LC/MS-MS, in DMSO-treated MV4-11 cells versus MV4-11 cells treated with 5 nM quizartinib or 5 μM WQ-2101 for 24 hours in complete RPMI, or serine/glycine-deprived RPMI. In all metabolomics experiments cell viability remained stable and was always >80%. All fold changes are relative to complete RPMI/DMSO control. All error bars are representative of ± s.d. and 4 independent biological replicates. Stars (*) are indicative of statistical comparisons between DMSO and treatment conditions as indicated. Hashes (#) are indicative of statistical comparisons within the same treatment condition, i.e., complete medium versus serine/glycine deprived medium for each condition. All statistical comparisons are performed by Student’s *t-*test. **B**. MV4-11, MOLM-13 and OCI-AML3 cells were treated for 24 hours with DMSO, 5 nM quizartinib or 5 μM WQ-2101 for 24 hours, prior to labelling with tritiated L-serine for 30 mins in serine/glycine-deprived RPMI medium. Cells were washed, lysed and tritiated serine uptake, measured in counts per minute (cpm), assessed by a scintillation counter. Error bars are representative of ± s.d. and 3 independent biological replicates. **** denotes *P* < 0.0001 by Student’s *t-*test. **C**. MV4-11 cells were treated with either DMSO or 3 nM quizartinib for 72 hours, and cocktails of purine nucleosides (adenine, guanine), pyrimidine nucleosides (cytosine, thymine, uridine) or all nucleosides were supplemented into medium for a final concentration of 150 μM at the time of seeding. 286 μM of L-serine was also supplemented into medium for a final concentration of 572 μM, and cell viability assessed via Annexin V flow cytometry. Statistical comparisons for each condition are performed relative to Quizartinib + Vehicle condition. Error bars are representative of ± s.d. and 3 independent biological replicates. ** denotes *P* < 0.01, *** *P* < 0.001 by Student’s *t-*test. **F**. MV4-11 cells were treated with either DMSO or 3 nM quizartinib for 48 hours, and metabolites relevant to PHGDH metabolism were supplemented into culture medium at the time of seeding. Final concentrations of metabolites supplemented were 500 μM pyruvate; 15 mM lactate; 150 μM hypoxanthine; 150 μM all nucleosides; and 286 μM of L-serine for a final concentration of 572 μM. Cell viability was determined by Annexin V flow cytometry, and all statistical comparisons for each condition performed relative to Quizartinib + Vehicle condition. Error bars are representative of ± s.d. and 3 independent biological replicates. * denotes *P* < 0.05, ** *P* < 0.01 by Student’s *t-*test. **E**. Analysis of a genetic signature encompassing genes involved in *de novo* serine synthesis, one-carbon/folate cycle, and rate-limiting *de novo* purine biosynthesis pathways (SSP-OC-Purine Signature). mRNA counts from the LAML-TCGA dataset were subset on FLT3-mutant (*n =* 41) and FLT3 wild-type (*n =* 100) samples, and the degree of correlation of the signature against *FLT3* mRNA expression in each subset was performed using the singscore scoring method (45).

The data to date strongly suggested that, in addition to serine metabolism, FLT3-ITD regulates the expression of genes important for one-carbon metabolism (**Figure 2**). Serine itself is a major one-carbon donor to the folate cycle, thereby facilitating the production of tetrahydrofolate (THF) intermediates required for nucleotide biosynthesis (23). Purine biosynthesis in particular has been shown to be heavily dependent on the mTORC1-ATF4 axis (43,46). Given that nucleotide imbalance is genotoxic to proliferating cells (47,48), it was possible that suppression of folate cycle activity and thus purine biosynthesis might contribute to the cytotoxicity associated with FLT3-ITD inhibition. Remarkably, supplementation of purine nucleosides considerably reversed the apoptotic effects of quizartinib treatment, whilst supplementation with pyrimidine nucleosides or a cocktail of purine and pyrimidine nucleosides had no capacity to rescue the cells (**Figure 5C**). Of note, doubling the exogenous supply of serine (to ∼572 μM) largely phenocopied the effects of purine supplementation, further supporting the contention that serine contributes to purine biosynthesis. As multiple metabolic processes are located upstream and downstream of PHGDH, MV4-11 culture media was therefore supplemented with metabolites relevant to PHGDH metabolism to determine the impact on cytotoxicity associated with FLT3-ITD inhibition (44). Supplementation with hypoxanthine, an adenine purine nucleobase derivative that undergoes conversion into the purine nucleotide inosine monophosphate (IMP), was sufficient to significantly reduce the apoptotic effects of quizartinib (**Figure 5D**).

To determine if these findings could be recapitulated in primary human samples, a genetic signature encompassing genes involved in *de novo* serine synthesis, the one-carbon/folate cycle, and key rate-limiting enzymes of the *de novo* purine synthesis pathway was devised as described by Manning and colleagues (43) (see also **Supplementary Materials and Methods**). As FLT3-mutant AMLs have been demonstrated to upregulate *FLT3* mRNA expression (49), stratification of the LAML-TCGA dataset by FLT3 mutation status showed a markedly greater, and significant positive correlation between this gene signature and *FLT3* mRNA expression in FLT3-mutant patient samples (**Figure 5E**). These data are consistent with the notion that FLT3-ITD inhibition reduces global serine availability and induces nucleotide insufficiency by both suppressing *de novo* serine synthesis pathway activity and serine uptake.

### Inhibition of *de novo* serine synthesis sensitizes FLT3-ITD-driven AMLs to cytarabine

Having demonstrated that PHGDH inhibition is a unique metabolic vulnerability in FLT3-ITD-mutant AML, strategies by which PHGDH inhibition could be exploited for the treatment of AML were explored. Cytarabine, the standard-of-care chemotherapeutic agent employed as front-line therapy in AML patients, is a deoxycytosine analogue that interferes with DNA synthesis by damaging DNA during the *S*-phase of replication.

Given the findings demonstrating purine nucleotide insufficiency upon FLT3-ITD inhibition (**Figure 5C**), it was possible that simultaneous treatment of AMLs with cytarabine and a PHGDH inhibitor would exacerbate DNA damage by restricting serine, and thus one-carbon units, for purine nucleotide synthesis and subsequent DNA repair processes. As FLT3-ITD-mutant leukemias displayed preferential sensitivity to PHGDH inhibition (**Figure 3E** and **Figure 3F**), it was likely that this combinatorial strategy would be particularly effective in this subset of AML.

Three FLT3-ITD positive AML cell lines; MV4-11, MOLM-13 and PL-21 were used for these studies. Of note, while MV4-11 and MOLM-13 cells display an *MLL*-translocated karyotype, PL-21 cells display a complex non-*MLL* karyotype (50). Exposure of all three cell lines to escalating concentrations of WQ-2101 in combination with cytarabine revealed synergistic interactions (highlighted in red), as determined using the SynergyFinder computational package and the Bliss synergy index (51) (**Figure 6A, Supplementary Figure 6A**). The combinatorial effect of cytarabine and WQ-2101 in MV4-11 cells was confirmed by Annexin-V staining (**Supplementary Figure 6B**). Consistent with the hypothesis that PHGDH inhibition would limit the availability of purine nucleotide species for DNA repair, WQ-2101 exacerbated cytarabine-induced phosphorylation of the DNA damage marker histone variant H2A.X in all three FLT3-ITD-mutant AML cell lines (**Figure 6B**).

**Figure 6.**
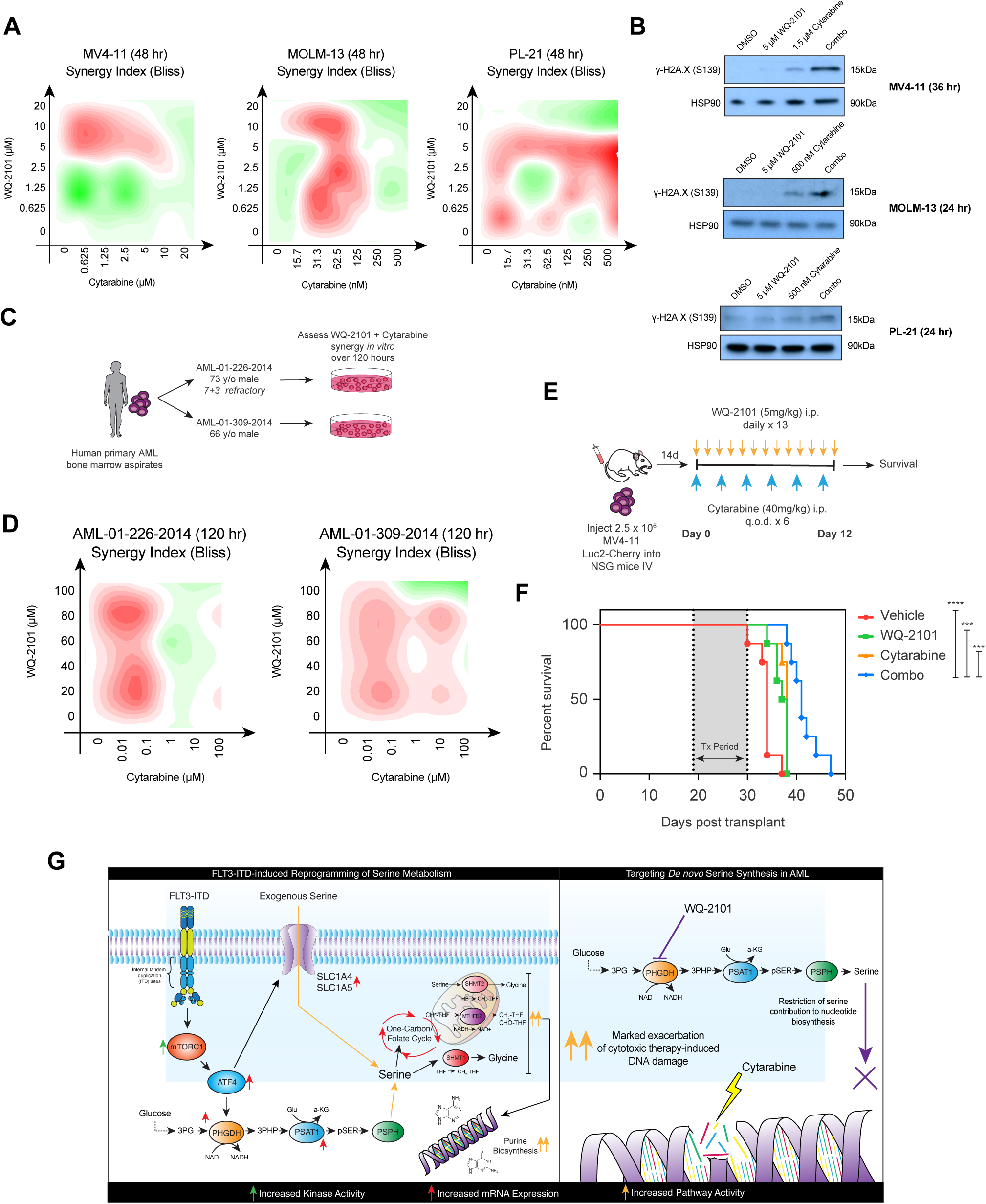
Inhibition of *de novo* serine synthesis sensitizes FLT3-ITD-driven AMLs to cytarabine. **A**. MV4-11, MOLM-13 and PL-21 cells were treated with escalating concentrations of WQ-2101 and cytarabine for 48 hours to determine the effect on viability, as determined by Cell Titer Glo. The presence of treatment synergy was determined using the SynergyFinder computational package and the Bliss synergy index and is denoted as regions of red in the graphs. The mean of three biological replicates was used to determine synergy. **B**. MV4-11 cells were treated with either DMSO, 5 μM WQ-2101, 1.5 μM cytarabine, or a combination of both, for 36 hours and DNA damage assessed via induction of γ-H2A.X, as determined by Western blot. MOLM-13 and PL-21 cells were treated with either DMSO, 5 μM WQ-2101, 500 nM cytarabine, or a combination of both, for 24 hours and DNA damage assessed as above. HSP90 served as the loading control in all experiments. **C**. Schematic representation of experiment testing combination therapy of WQ-2101 and cytarabine *in vitro* in primary patient AML bone marrow aspirates. **D**. Cells from two independent primary AMLs, AML-01-226-2014 and AML-01-309-2014, were treated with escalating doses of WQ-2101 and cytarabine for 120 hours to determine the effect on cell viability, as determined by Sytox Blue flow cytometry. The presence of treatment synergy was determined using the SynergyFinder computational package and the Bliss synergy index and is denoted as regions of red in the graphs. Background death of AML-01-226-2014 and AML-01-309-2014 upon thaw was 10.4% and 15.5% respectively. Data is indicative of one experiment. **E**. Schematic diagram of *in vivo* combination therapy experimental protocol, using an aggressive human MV4-11 PDX disseminated model in NSG mice. **F**. Kaplan-Meier survival analysis of MV4-11 disseminated PDX model in NSG mice. Mice were treated with Vehicle, 5 mg/kg WQ-2101 daily, 40 mg/kg cytarabine every second day, or a combination of both for 12 days. All statistical comparisons are combination therapy relative to all other treatment conditions as indicated. *** denotes *P* < 0.001, **** *P* < 0.0001 as determined by Mantel-Cox test, with *n =* 8 mice per condition. **G**. Schematic outline of this study. FLT3-ITD transcriptionally modulates *de novo* serine synthesis and serine uptake, as well as downstream one-carbon/folate cycle activity. This results in heightened purine biosynthesis. Thus, pharmacological inhibition of PHGDH sensitizes AML cells to standard-of-care cytarabine-induced DNA damage by exacerbating the DNA damage response.

To determine if this combinatorial strategy has potential clinical implications, two FLT3-ITD-mutant primary AML bone marrow aspirates termed AML-01-226-2014 and AML-01-309-2014 were sourced (**Figure 6C** – clinical data available in **Supplementary Figure 6C**). Of note, AML-01-226-2014 was refractory to standard-of-care 7+3 therapy with cytarabine and daunorubicin. Consistent with data shown using FLT3-ITD-mutant human AML cell lines, a combinatorial effect between cytarabine and WQ-2101 was evident in both primary AML samples (**Figure 6D, Supplementary Figure 6D**).

Finally, to determine if this combinatorial strategy has *in vivo* efficacy, mice bearing disseminated MV4-11 cells were treated with single agent cytarabine, WQ-2101, or a combination of both (**Figure 6E**). Single agent WQ-2101 and cytarabine provided limited survival advantage to leukemic mice relative to vehicle treatment, while the combinatorial arm displayed significant survival advantage relative to vehicle and both WQ-2101 and cytarabine single agent arms (**Figure 6F**). These *in vitro* data in human AML cell lines and primary patient specimens, in conjunction with evidence of *in vivo* activity in an aggressive disseminated model of AML, provide a strong rationale for targeting dysregulated serine synthesis in conjunction with standard of care chemotherapy in FLT3-ITD mutant AML (**Figure 6G**).

## DISCUSSION

The recent approval of the first generation pan-receptor tyrosine kinase (RTK) inhibitor midostaurin as frontline therapy for patients with FLT3-ITD mutations has resulted in significant interest in targeting FLT3 in the context of AML (7). Although FLT3 inhibitor monotherapy is promising and initially yielded encouraging results, the development of acquired resistance has reduced the efficacy of this approach. Here, we developed a mouse model of *MLL*-rearranged, FLT3-ITD AML to reveal FLT3-ITD-regulated pathways important for sustainable AML growth and to identify new therapeutic targets. We discovered that serine metabolism was necessary for the proliferation and survival of MLL-rearranged FLT3-ITD leukemias, thus revealing a targetable vulnerability in this subset of AML that could be exploited using small molecule inhibitors of the *de novo* serine synthesis pathway.

The oncogenic functions of MLL-AF9 have been well documented with numerous studies detailing the role of this fusion protein in de-regulating the epigenetic and transcriptional landscape to drive leukemogenesis (32). A wide variety of mouse models of *MLL*-rearranged AML have thus been established (52) with co-introduction of mutant *Ras* (31) or *FLT3-ITD* (28) to accelerate leukaemic development. We and others have previously shown that genetic de-induction of *MLL-AF9* in these models results in terminal differentiation through the activity of transcriptional regulators such as *Myb* (33) and the *Id2*/E-protein axis (31). While the biological consequences of inducible *Nras* depletion in AML has been studied (53,54), to the best of our knowledge, the work described herein is the first study to address the importance of FLT3-ITD in the maintenance of MLL-AF9-driven AML.

A key finding from this study is that FLT3-ITD modulates serine metabolism, by upregulating mRNA expression of genes such as *PHGDH* and *PSAT1*, and serine uptake through enhanced expression of the neutral amino acid transporters *SLC1A4* and *SLC1A5*. Serine is a major source of one-carbon units for the tetrahydrofolate cycle, which provides metabolic outputs important for maintenance of adenosine triphosphate (ATP), and *S*-adenosyl methionine (SAM) pools, as well as downstream biosynthesis of purine and pyrimidine species (24). While most cellular serine requirements are met via uptake of exogenous serine, genomic and non-genomic events can divert glucose-derived carbons to the *de novo* serine synthesis pathway. Indeed, genomic analyses in primary breast cancer, melanoma and prostate cancers identified copy number amplification in enzymes of the *de novo* serine synthesis pathway, most notably phosphoglycerate dehydrogenase (*PHGDH)*, the first and only rate-limiting enzyme of this pathway (55-58). Importantly, transcriptional mechanisms that mediate activation of the *de novo* serine synthesis pathway have also been discovered. *ATF4*, a transcription factor that responds to cellular stressors such as amino acid starvation and hypoxia, and was identified as central to FLT3-ITD-mediated regulation of the serine synthesis pathway in this study, has been previously demonstrated to activate key enzymes of the *de novo* serine synthesis pathway in other disease contexts (40,59). Other transcription factors such as *MYC* (60,61) and *NRF2* via *ATF4* (62) have also been demonstrated to induce *de novo* serine synthesis pathway activity. Interestingly, *RAS* pathway activation has been shown to enhance serine biosynthesis. *KRAS* activation and concurrent loss of the tumour suppressor gene *LKB1* enhances proliferative capacity in pancreatic cancer models, primary via stimulation of serine biosynthesis (63). A second study demonstrated lack of sensitivity to restriction of dietary serine in pancreatic and intestinal tumours, due to the inherent ability of *KRAS* mutations to activate *de novo* serine synthesis (64).

The data from our study and others shows that the mTORC1-ATF4 axis is pivotal to the induction of serine biosynthetic pathways. The link between FLT3-ITD and mTORC1 has been previously defined (41) and mTORC1 subsequently controls ATF4 via two distinct methods: (i) by regulating the stability of ATF4 mRNA transcripts; (ii) via regulation of ATF4 translation through three upstream open reading frames (uORFs) in the 5’ UTR of ATF4 (42). In the context of FLT3-ITD mutations in AML, we provide a model whereby mTORC1 activates ATF4 to drive transcriptional regulation of both the *de novo* serine synthesis pathway and several key transmembrane transporters associated with serine/glycine import into the cell. Previous studies have demonstrated that FLT3-ITD-driven AMLs are preferentially sensitive to mTOR/PI3K inhibitors (65-68), and it is possible that at least part of their efficacy can be attributed to suppression of the mTORC1/ATF4 axis, and disruption of downstream metabolic pathways such as the *de novo* serine synthesis pathway.

There has been considerable interest in the development of small molecule inhibitors to target the *de novo* serine synthesis pathway, and small molecule PHGDH inhibitors have been developed, primarily investigated in the context of solid malignancies such as breast and prostate cancers (36,69-72). Our molecular analysis of FLT3-ITD-driven AML provided a strong rationale to apply the PHGDH inhibitor WQ-2101 in our leukemia models to exploit the dependency of these tumours on sustained serine biosynthesis. Why tumours, including AMLs, display sensitivity to PHGDH inhibition even in the presence of exogenous serine is an important question when considering the efficacy of these compounds. One possible explanation is that *de novo*-generated serine may ensure availability of one-carbon units for nucleotide biosynthesis by reducing cytoplasmic *SHMT1* activity (70). Inhibition of PHGDH (and, therefore, activation of *SHMT1*) even in contexts in which exogenous serine is abundant, reduces the incorporation of one-carbon units into nucleotides which may result in the inability of exogenous serine to rescue the loss of PHGDH. Another study investigating WQ-2101 discovered that PHGDH inhibition affects nucleotide synthesis in a manner that is independent of serine utilization, by disrupting central carbon metabolism, the pentose-phosphate pathway and TCA cycle (44). This downstream effect on nucleotide metabolism was explored in this study. As we have demonstrated, FLT3-ITD inhibition transcriptionally suppresses *de novo* serine synthesis and one-carbon cycle activity, thereby triggering downstream purine insufficiency. The *de novo* serine synthesis pathway donates carbon units to the folate cycle, which produces cytosolic one-carbon formyl units that are critical for purine ring assembly (43,46). Inhibition of purine metabolism has long been explored as a therapeutic modality in leukaemia (73) and more recently, several groups have investigated the therapeutic implications of perturbing enzymes in the *de novo* purine synthesis pathway (such as *PAICS*) in the context of AML (74). We showed that inhibition of PHGDH sensitized FLT3-ITD AML to cytarabine and posit that this is through enhanced chemotherapy-induced DNA damage. Our biochemical studies demonstrating greatly enhanced gH2AX signal following combined cytarabine and WQ-2101 treatment is consistent with this hypothesis.

Collectively, our study reveals new mechanistic insights into FLT3-ITD-induced metabolic reprogramming events in AML, and reveals serine metabolism as a unique therapeutic vulnerability in this subset of AML. Our work also highlights how the therapeutic utility of PHGDH inhibition can be maximised by combining *de novo* serine synthesis pathway inhibitors with standard of care chemotherapy agents such as cytarabine to exacerbate DNA damage responses and provide therapeutic benefit in pre-clinical AML models.

## METHODS

### Cell lines and cell culture

MLL-AF9/iFLT3-ITD murine cells were generated via retroviral transduction of E13.5-day foetal liver cells as described below, and cultured in Anne Kelso Modified Dulbecco’s Modified Eagle Medium (AK-DMEM – Low glucose DMEM (Invitrogen Life Technologies), 4 g/L glucose, 36 mg/mL folic acid, 116 mg/L L-arginine HCl, 216 mg/L L-glutamine, 2 g/L NaHCO_3_ and 10 mM HEPES), supplemented with 20% foetal bovine serum (FBS) (Invitrogen Life Technologies), 1% penicillin/streptomycin (PenStrep) (Invitrogen Life Technologies) and 1% L-asparagine (Sigma-Aldrich) at 37ºC and 10% CO_2_. Cells at baseline were cultured in 1μg/mL doxycycline (Sigma-Aldrich) to maintain transgene expression. MV4-11, Kasumi-1, HL-60 and HEK-293T/17 cells were purchased from the ATCC. MOLM-13, OCI-AML3 and PL-21 cells were purchased from the DSMZ. MV4-11, MOLM-13 and PL-21 cells were cultured in RPMI-1640 (Gibco) supplemented with 10% FBS, 1% PenStrep and 1% Glutamax (Thermo Fisher). Kasumi-1, HL-60 and OCI-AML3 cells were cultured in the same conditions, with the exception of 20% FBS content. HEK-293T cells were cultured in DMEM (Gibco), supplemented with 10% FBS, 1% PenStrep, and 1% Glutamax. For serine/glycine deprivation experiments, we synthesised RPMI-1640 and omitted inclusion of serine and glycine during preparation of medium. Exact composition of media is supplied in **Supplementary Materials and Methods**. All human lines were cultured at 37ºC and at 5% CO_2_. All cells used in this study were tested *mycoplasma* negative via PCR validation. All cell lines were validated via short-tandem repeat (*STR*) profiling prior to use.

### Compounds and chemicals

Quizartinib, WQ-2101 and Everolimus small molecule inhibitors were purchased from Selleckchem, and reconstituted in DMSO for *in vitro* use. Fresh cytarabine in saline (AraC) was acquired from the Peter MacCallum Cancer Centre cytotoxic suite. *In vivo* quantities of WQ-2101 were purchased from Glixx Labs and reconstituted in 10% DMSO, 20% Kolliphor HS-15 and 70% sterile water supplemented with 0.1% Tween 80 as previously described (36). For *in vivo* dosing, WQ-2101 was reconstituted daily. AraC was diluted in PBS as required.

### Retroviral/lentiviral transductions and plasmid cloning

MLL-AF9/iFLT3-ITD murine cell lines were generated via retroviral transduction of E13.5 foetal liver cells from a mouse endogenously expressing the CAG-rtTA3 transgene as described in **Supplementary Materials and Methods**. For lentiviral transductions for CRISPR-Cas9-mediated depletion of *PHGDH*, and lentiviral transductions for shRNA-mediated silencing of *ATF4*, see **Supplementary Materials and Methods**.

### Western immunoblotting

Whole-cell protein lysate was obtained by suspension of cells in Laemmli lysis buffer at room temperature for 10 minutes, followed by boiling of samples at 95ºC for 10 minutes. Protein lysates were separated by SDS-PAGE and transferred to polyvinylidene difluoride (PVDF) membranes. The membranes were blocked in 5% skim milk and incubated with primary antibody overnight. Membranes were subsequently washed in TBS-T and incubated with horseradish peroxidase (HRP)-conjugated secondary antibody as appropriate. Blots were developed using enhanced chemiluminescence (ECL) and ECL Prime (GE Healthcare). For blots probing for phosphorylated and total targets, a master mix of each lysate was prepared and an equal volume of protein run on separate blots, with each blot probed with appropriate primary antibody. Loading controls were developed with representative examples shown in each figure. For a complete list of primary antibodies and secondary antibodies, see **Supplementary Materials and Methods**.

### Cell death assays

For assessment of cell viability via Annexin V staining, cell suspension was stained with 1:100 dilution Annexin V conjugated to an APC fluorophore (BD Pharmingen, Cat #550475) in binding buffer (10 mM HEPES, 5 mM CaCl_2_, 140 mM NaCl_2_) prior to flow cytometry analysis. For assessment of cell viability in response to WQ-2101, 8,000 AML cells were plated into 384-well plates and exposed to escalating doses of WQ-2101 for 48 hours. At endpoint, 1:2 dilution Cell Titer Glo (CTG) was added to each well and cells lysed for 10 minutes prior to assessment of absorbance with a Cytation 3 spectrophotometer (BioTek).

### Competition assays and assessment of cell proliferation

For competition assays, Tet-inducible sgRNAs were activated by addition of doxycycline to cell medium 5 days prior to the assay. 50:50 mixes of parental and GFP-positive sgRNA containing cells were then cultured in 6-well plates and the ratio of parental to GFP-positive cells assessed via flow cytometry over 14 days. For determination of proliferation in the presence/absence of serine and WQ-2101, 50,000 cells were seeded in 6-well plates and cultured for 3 or 9 days respectively. Cells were counted using a cellometer, and viability determined by trypan-exclusion assay. For Cell Trace Violet (CTV) analysis of cell proliferation in response to WQ-2101, 1 × 10^7^ cells were washed in PBS, then stained for 20 minutes at 37ºC with agitation with Cell Trace Violet label (Thermo Fisher). A narrow peak of cells was then sorted using a FACSAria Fusion 5 (BD) to synchronise cell cycle phase and enable defined peak analysis. Cells were then cultured for 72 hours in 3 μM WQ-2101 and proliferation assessed via flow cytometry.

### QuantSeq 3’RNA-sequencing

Details of 3’RNA-sequencing RNA preparation, sequencing and analysis pipelines can be found in **Supplementary Materials and Methods**.

### Chromatin immunoprecipitation sequencing

Details of ChIP-sequencing protocols, library preparation, sequencing and analysis pipelines can be found in **Supplementary Materials and Methods**.

### Metabolomics

MV4-11 cells were treated as described, harvested, and processed for steady-state metabolomics analysis as described in **Supplementary Materials and Methods**.

### Metabolic rescue experiments

Detailed methods for serine rescue experiments in sgPHGDH cells and metabolite rescue experiments in response to FLT3-ITD inhibition with quizartinib, can be found in **Supplementary Materials and Methods**.

### Tritiated serine labelling

Detailed methods for assessment of exogenous serine uptake post FLT3-ITD and PHGDH inhibition with quizartinib and WQ-2101 can be found in **Supplementary Materials and Methods**.

### Combination therapy analysis *in vitro* and *in vivo*

Experimental protocols and analysis for combination therapy with WQ-2101 and cytarabine in AML cell lines, primary patient samples and *in vivo*, can be found in **Supplementary Materials and Methods**.

### Statistical analyses

GraphPad Prism v8.0 and R version 3.6.1 software were used for statistical analysis. Statistical tests performed for each experiment are highlighted in the figure legends. Unless otherwise specified, error bars are indicative of plus/minus standard deviation (± s.d.) and threshold for significance was *P* < 0.05.

### Data availability

RNA-seq and ChIP-seq data was deposited in the Gene Expression Omnibus (GEO). Outputs from RNA-seq DGE analysis and ChIP-seq peak calling are including in the **Supplementary Data Tables**. Any other relevant raw data is available from the corresponding authors at reasonable request.

## Acknowledgements

We acknowledge support from the Peter MacCallum Cancer Centre Foundation, Australian Cancer Research Foundation and Metabolomics Australia at the University of Melbourne (MA at UoM).

## REFERENCES

1. Patel JP, Gonen M, Figueroa ME, Fernandez H, Sun Z, Racevskis J, et al. Prognostic relevance of integrated genetic profiling in acute myeloid leukemia. N Engl J Med 2012;366(12):1079–89 doi 10.1056/NEJMoa1112304.

2. Papaemmanuil E, Gerstung M, Bullinger L, Gaidzik VI, Paschka P, Roberts ND, et al. Genomic Classification and Prognosis in Acute Myeloid Leukemia. 2016;374(23):2209–21 doi 10.1056/NEJMoa1516192.

3. Metzeler KH, Herold T, Rothenberg-Thurley M, Amler S, Sauerland MC, Gorlich D, et al. Spectrum and prognostic relevance of driver gene mutations in acute myeloid leukemia. Blood 2016;128(5):686–98 doi 10.1182/blood-2016-01-693879.

4. Tyner JW, Tognon CE, Bottomly D, Wilmot B, Kurtz SE, Savage SL, et al. Functional genomic landscape of acute myeloid leukaemia. Nature 2018;562(7728):526–31 doi 10.1038/s41586-018-0623-z.

5. Talati C, Sweet K. Recently approved therapies in acute myeloid leukemia: A complex treatment landscape. Leuk Res 2018;73:58–66 doi 10.1016/j.leukres.2018.09.001.

6. Short NJ, Rytting ME, Cortes JE. Acute myeloid leukaemia. Lancet 2018;392(10147):593–606 doi 10.1016/S0140-6736(18)31041-9.

7. Stone RM, Mandrekar SJ, Sanford BL, Laumann K, Geyer S, Bloomfield CD, et al. Midostaurin plus Chemotherapy for Acute Myeloid Leukemia with a FLT3 Mutation. N Engl J Med 2017;377(5):454–64 doi 10.1056/NEJMoa1614359.

8. Smith CC, Wang Q, Chin CS, Salerno S, Damon LE, Levis MJ, et al. Validation of ITD mutations in FLT3 as a therapeutic target in human acute myeloid leukaemia. Nature 2012;485(7397):260–3 doi 10.1038/nature11016.

9. Kenins L, Gill JW, Hollander GA, Wodnar-Filipowicz A. Flt3 ligand-receptor interaction is important for maintenance of early thymic progenitor numbers in steady-state thymopoiesis. Eur J Immunol 2010;40(1):81–90 doi 10.1002/eji.200839213.

10. Zhang S, Fukuda S, Lee Y, Hangoc G, Cooper S, Spolski R, et al. Essential role of signal transducer and activator of transcription (Stat)5a but not Stat5b for Flt3-dependent signaling. J Exp Med 2000;192(5):719–28 doi 10.1084/jem.192.5.719.

11. Choudhary C, Brandts C, Schwable J, Tickenbrock L, Sargin B, Ueker A, et al. Activation mechanisms of STAT5 by oncogenic Flt3-ITD. Blood 2007;110(1):370–4 doi 10.1182/blood-2006-05-024018.

12. Lindblad O, Cordero E, Puissant A, Macaulay L, Ramos A, Kabir NN, et al. Aberrant activation of the PI3K/mTOR pathway promotes resistance to sorafenib in AML. Oncogene 2016;35(39):5119–31 doi 10.1038/onc.2016.41.

13. Takahashi S. Inhibition of the MEK/MAPK signal transduction pathway strongly impairs the growth of Flt3-ITD cells. Am J Hematol 2006;81(2):154–5 doi 10.1002/ajh.20520.

14. McMahon CM, Ferng T, Canaani J, Wang ES, Morrissette JJD, Eastburn DJ, et al. Clonal Selection with RAS Pathway Activation Mediates Secondary Clinical Resistance to Selective FLT3 Inhibition in Acute Myeloid Leukemia. Cancer Discov 2019;9(8):1050–63 doi 10.1158/2159-8290.CD-18-1453.

15. Vander Heiden MG, DeBerardinis RJ. Understanding the Intersections between Metabolism and Cancer Biology. Cell 2017;168(4):657–69 doi 10.1016/j.cell.2016.12.039.

16. Luengo A, Gui DY, Vander Heiden MG. Targeting Metabolism for Cancer Therapy. Cell Chemical Biology 2017;24(9):1161–80 doi https://doi.org/10.1016/j.chembiol.2017.08.028.

17. Brown KK, Spinelli JB, Asara JM, Toker A. Adaptive Reprogramming of De Novo Pyrimidine Synthesis Is a Metabolic Vulnerability in Triple-Negative Breast Cancer. Cancer Discov 2017;7(4):391–9 doi 10.1158/2159-8290.Cd-16-0611.

18. Poulain L, Sujobert P, Zylbersztejn F, Barreau S, Stuani L, Lambert M, et al. High mTORC1 activity drives glycolysis addiction and sensitivity to G6PD inhibition in acute myeloid leukemia cells. Leukemia 2017;31(11):2326–35 doi 10.1038/leu.2017.81.

19. Ju HQ, Zhan G, Huang A, Sun Y, Wen S, Yang J, et al. ITD mutation in FLT3 tyrosine kinase promotes Warburg effect and renders therapeutic sensitivity to glycolytic inhibition. Leukemia 2017;31(10):2143–50 doi 10.1038/leu.2017.45.

20. Gallipoli P, Giotopoulos G, Tzelepis K, Costa ASH, Vohra S, Medina-Perez P, et al. Glutaminolysis is a metabolic dependency in FLT3(ITD) acute myeloid leukemia unmasked by FLT3 tyrosine kinase inhibition. Blood 2018;131(15):1639–53 doi 10.1182/blood-2017-12-820035.

21. Gregory MA, Nemkov T, Reisz JA, Zaberezhnyy V, Hansen KC, D’Alessandro A, et al. Glutaminase inhibition improves FLT3 inhibitor therapy for acute myeloid leukemia. Exp Hematol 2018;58:52–8 doi 10.1016/j.exphem.2017.09.007.

22. Ducker GS, Rabinowitz JD. One-Carbon Metabolism in Health and Disease. Cell Metab 2017;25(1):27–42 doi 10.1016/j.cmet.2016.08.009.

23. Yang M, Vousden KH. Serine and one-carbon metabolism in cancer. Nat Rev Cancer 2016;16(10):650–62 doi 10.1038/nrc.2016.81.

24. Newman AC, Maddocks ODK. Serine and Functional Metabolites in Cancer. Trends Cell Biol 2017;27(9):645–57 doi 10.1016/j.tcb.2017.05.001.

25. Lee BH, Williams IR, Anastasiadou E, Boulton CL, Joseph SW, Amaral SM, et al. FLT3 internal tandem duplication mutations induce myeloproliferative or lymphoid disease in a transgenic mouse model. Oncogene 2005;24(53):7882–92 doi 10.1038/sj.onc.1208933.

26. Shih LY, Huang CF, Wang PN, Wu JH, Lin TL, Dunn P, et al. Acquisition of FLT3 or N-ras mutations is frequently associated with progression of myelodysplastic syndrome to acute myeloid leukemia. Leukemia 2004;18(3):466–75 doi 10.1038/sj.leu.2403274.

27. Thiede C, Steudel C, Mohr B, Schaich M, Schakel U, Platzbecker U, et al. Analysis of FLT3-activating mutations in 979 patients with acute myelogenous leukemia: association with FAB subtypes and identification of subgroups with poor prognosis. Blood 2002;99(12):4326–35 doi 10.1182/blood.v99.12.4326.

28. Stubbs MC, Kim YM, Krivtsov AV, Wright RD, Feng Z, Agarwal J, et al. MLL-AF9 and FLT3 cooperation in acute myelogenous leukemia: development of a model for rapid therapeutic assessment. Leukemia 2008;22(1):66–77 doi 10.1038/sj.leu.2404951.

29. Moullan N, Mouchiroud L, Wang X, Ryu D, Williams EG, Mottis A, et al. Tetracyclines Disturb Mitochondrial Function across Eukaryotic Models: A Call for Caution in Biomedical Research. Cell Rep 2015;10(10):1681–91 doi 10.1016/j.celrep.2015.02.034.

30. Ahler E, Sullivan WJ, Cass A, Braas D, York AG, Bensinger SJ, et al. Doxycycline alters metabolism and proliferation of human cell lines. PLoS One 2013;8(5):e64561. doi 10.1371/journal.pone.0064561.

31. Ghisi M, Kats L, Masson F, Li J, Kratina T, Vidacs E, et al. Id2 and E Proteins Orchestrate the Initiation and Maintenance of MLL-Rearranged Acute Myeloid Leukemia. Cancer Cell 2016;30(1):59–74 doi 10.1016/j.ccell.2016.05.019.

32. Winters AC, Bernt KM. MLL-Rearranged Leukemias-An Update on Science and Clinical Approaches. Front Pediatr 2017;5:4- doi 10.3389/fped.2017.00004.

33. Zuber J, Rappaport AR, Luo W, Wang E, Chen C, Vaseva AV, et al. An integrated approach to dissecting oncogene addiction implicates a Myb-coordinated self-renewal program as essential for leukemia maintenance. Genes Dev 2011;25(15):1628–40 doi 10.1101/gad.17269211.

34. Labuschagne CF, van den Broek NJ, Mackay GM, Vousden KH, Maddocks OD. Serine, but not glycine, supports one-carbon metabolism and proliferation of cancer cells. Cell Rep 2014;7(4):1248–58 doi 10.1016/j.celrep.2014.04.045.

35. Cantor JR, Abu-Remaileh M, Kanarek N, Freinkman E, Gao X, Louissaint A, Jr., et al. Physiologic Medium Rewires Cellular Metabolism and Reveals Uric Acid as an Endogenous Inhibitor of UMP Synthase. Cell 2017;169(2):258–72.e17 doi 10.1016/j.cell.2017.03.023.

36. Wang Q, Liberti MV, Liu P, Deng X, Liu Y, Locasale JW, et al. Rational Design of Selective Allosteric Inhibitors of PHGDH and Serine Synthesis with Anti-tumor Activity. Cell Chem Biol 2017;24(1):55–65 doi 10.1016/j.chembiol.2016.11.013.

37. Chao Q, Sprankle KG, Grotzfeld RM, Lai AG, Carter TA, Velasco AM, et al. Identification of N-(5-tert-Butyl-isoxazol-3-yl)-N′-{4-[7-(2-morpholin-4-yl-ethoxy)imidazo[2,1-b][1,3]benzothiazol-2-yl]phenyl}urea Dihydrochloride (AC220), a Uniquely Potent, Selective, and Efficacious FMS-Like Tyrosine Kinase-3 (FLT3) Inhibitor. Journal of Medicinal Chemistry 2009;52(23):7808–16 doi 10.1021/jm9007533.

38. Zarrinkar PP, Gunawardane RN, Cramer MD, Gardner MF, Brigham D, Belli B, et al. AC220 is a uniquely potent and selective inhibitor of FLT3 for the treatment of acute myeloid leukemia (AML). Blood 2009;114(14):2984–92 doi 10.1182/blood-2009-05-222034.

39. Imrichova H, Hulselmans G, Atak ZK, Potier D, Aerts S. i-cisTarget 2015 update: generalized cis-regulatory enrichment analysis in human, mouse and fly. Nucleic Acids Res 2015;43(W1):W57–64 doi 10.1093/nar/gkv395.

40. Ye J, Mancuso A, Tong X, Ward PS, Fan J, Rabinowitz JD, et al. Pyruvate kinase M2 promotes de novo serine synthesis to sustain mTORC1 activity and cell proliferation. Proc Natl Acad Sci U S A 2012;109(18):6904–9 doi 10.1073/pnas.1204176109.

41. Chen W, Drakos E, Grammatikakis I, Schlette EJ, Li J, Leventaki V, et al. mTOR signaling is activated by FLT3 kinase and promotes survival of FLT3-mutated acute myeloid leukemia cells. Mol Cancer 2010;9:292 doi 10.1186/1476-4598-9-292.

42. Park Y, Reyna-Neyra A, Philippe L, Thoreen CC. mTORC1 Balances Cellular Amino Acid Supply with Demand for Protein Synthesis through Post-transcriptional Control of ATF4. Cell Rep 2017;19(6):1083–90 doi 10.1016/j.celrep.2017.04.042.

43. Ben-Sahra I, Hoxhaj G, Ricoult SJH, Asara JM, Manning BD. mTORC1 induces purine synthesis through control of the mitochondrial tetrahydrofolate cycle. Science 2016;351(6274):728–33 doi 10.1126/science.aad0489.

44. Reid MA, Allen AE, Liu S, Liberti MV, Liu P, Liu X, et al. Serine synthesis through PHGDH coordinates nucleotide levels by maintaining central carbon metabolism. Nat Commun 2018;9(1):5442 doi 10.1038/s41467-018-07868-6.

45. Foroutan M, Bhuva DD, Lyu R, Horan K, Cursons J, Davis MJ. Single sample scoring of molecular phenotypes. BMC Bioinformatics 2018;19(1):404 doi 10.1186/s12859-018-2435-4.

46. Villa E, Ali ES, Sahu U, Ben-Sahra I. Cancer Cells Tune the Signaling Pathways to Empower de Novo Synthesis of Nucleotides. Cancers (Basel) 2019;11(5) doi 10.3390/cancers11050688.

47. Kunz BA. Mutagenesis and deoxyribonucleotide pool imbalance. Mutat Res 1988;200(1-2):133–47 doi 10.1016/0027-5107(88)90076-0.

48. Kunz BA, Kohalmi SE, Kunkel TA, Mathews CK, McIntosh EM, Reidy JA. International Commission for Protection Against Environmental Mutagens and Carcinogens. Deoxyribonucleoside triphosphate levels: a critical factor in the maintenance of genetic stability. Mutat Res 1994;318(1):1–64 doi 10.1016/0165-1110(94)90006-x.

49. Cheng J, Qu L, Wang J, Cheng L, Wang Y. High expression of FLT3 is a risk factor in leukemia. Mol Med Rep 2018;17(2):2885–92 doi 10.3892/mmr.2017.8232.

50. Kubonishi I, Machida K, Niiya K, Sonobe H, Ohtsuki Y, Iwata K, et al. Establishment of a new peroxidase-positive human myeloid cell line, PL-21. Blood 1984;63(2):254–9.

51. Ianevski A, He L, Aittokallio T, Tang J. SynergyFinder: a web application for analyzing drug combination dose-response matrix data. Bioinformatics 2017;33(15):2413–5 doi 10.1093/bioinformatics/btx162.

52. Milne TA. Mouse models of MLL leukemia: recapitulating the human disease. Blood 2017;129(16):2217–23 doi 10.1182/blood-2016-10-691428.

53. Kim WI, Matise I, Diers MD, Largaespada DA. RAS oncogene suppression induces apoptosis followed by more differentiated and less myelosuppressive disease upon relapse of acute myeloid leukemia. Blood 2009;113(5):1086–96 doi 10.1182/blood-2008-01-132316. Epub 2008 Oct 24. papers3://publication/doi/10.1182/blood-2008-01-132316.

54. Sachs Z, LaRue RS, Nguyen HT, Sachs K, Noble KE, Mohd Hassan NA, et al. NRASG12V oncogene facilitates self-renewal in a murine model of acute myelogenous leukemia. Blood 2014;124(22):3274–83 doi 10.1182/blood-2013-08-521708. Epub 2014 Oct 14. papers3://publication/doi/10.1182/blood-2013-08-521708.

55. Possemato R, Marks KM, Shaul YD, Pacold ME, Kim D, Birsoy K, et al. Functional genomics reveal that the serine synthesis pathway is essential in breast cancer. Nature 2011;476(7360):346–50 doi 10.1038/nature10350.

56. Mehrmohamadi M, Liu X, Shestov AA, Locasale JW. Characterization of the usage of the serine metabolic network in human cancer. Cell Rep 2014;9(4):1507–19 doi 10.1016/j.celrep.2014.10.026.

57. Locasale JW, Grassian AR, Melman T, Lyssiotis CA, Mattaini KR, Bass AJ, et al. Phosphoglycerate dehydrogenase diverts glycolytic flux and contributes to oncogenesis. Nat Genet 2011;43(9):869–74 doi 10.1038/ng.890.

58. Beroukhim R, Mermel CH, Porter D, Wei G, Raychaudhuri S, Donovan J, et al. The landscape of somatic copy-number alteration across human cancers. Nature 2010;463(7283):899–905 doi 10.1038/nature08822.

59. Adams CM. Role of the transcription factor ATF4 in the anabolic actions of insulin and the anti-anabolic actions of glucocorticoids. J Biol Chem 2007;282(23):16744–53 doi 10.1074/jbc.M610510200.

60. Nilsson LM, Forshell TZ, Rimpi S, Kreutzer C, Pretsch W, Bornkamm GW, et al. Mouse genetics suggests cell-context dependency for Myc-regulated metabolic enzymes during tumorigenesis. PLoS Genet 2012;8(3):e1002573.doi 10.1371/journal.pgen.1002573.

61. Sun L, Song L, Wan Q, Wu G, Li X, Wang Y, et al. cMyc-mediated activation of serine biosynthesis pathway is critical for cancer progression under nutrient deprivation conditions. Cell Res 2015;25(4):429–44 doi 10.1038/cr.2015.33.

62. DeNicola GM, Chen PH, Mullarky E, Sudderth JA, Hu Z, Wu D, et al. NRF2 regulates serine biosynthesis in non-small cell lung cancer. Nat Genet 2015;47(12):1475–81 doi 10.1038/ng.3421.

63. Kottakis F, Nicolay BN, Roumane A, Karnik R, Gu H, Nagle JM, et al. LKB1 loss links serine metabolism to DNA methylation and tumorigenesis. Nature 2016;539(7629):390–5 doi 10.1038/nature20132.

64. Maddocks ODK, Athineos D, Cheung EC, Lee P, Zhang T, van den Broek NJF, et al. Modulating the therapeutic response of tumours to dietary serine and glycine starvation. Nature 2017;544(7650):372–6 doi 10.1038/nature22056.

65. Barrett D, Brown VI, Grupp SA, Teachey DT. Targeting the PI3K/AKT/mTOR signaling axis in children with hematologic malignancies. Paediatr Drugs 2012;14(5):299–316 doi 10.2165/11594740-000000000-00000.

66. Martelli AM, Evangelisti C, Chappell W, Abrams SL, Basecke J, Stivala F, et al. Targeting the translational apparatus to improve leukemia therapy: roles of the PI3K/PTEN/Akt/mTOR pathway. Leukemia 2011;25(7):1064–79 doi 10.1038/leu.2011.46.

67. Nogami A, Oshikawa G, Okada K, Fukutake S, Umezawa Y, Nagao T, et al. FLT3-ITD confers resistance to the PI3K/Akt pathway inhibitors by protecting the mTOR/4EBP1/Mcl-1 pathway through STAT5 activation in acute myeloid leukemia. Oncotarget 2015;6(11):9189–205 doi 10.18632/oncotarget.3279.

68. Chapuis N, Tamburini J, Green AS, Vignon C, Bardet V, Neyret A, et al. Dual inhibition of PI3K and mTORC1/2 signaling by NVP-BEZ235 as a new therapeutic strategy for acute myeloid leukemia. Clin Cancer Res 2010;16(22):5424–35 doi 10.1158/1078-0432.Ccr-10-1102.

69. Mullarky E, Lairson LL, Cantley LC, Lyssiotis CA. A novel small-molecule inhibitor of 3-phosphoglycerate dehydrogenase. Mol Cell Oncol 2016;3(4):e1164280.doi 10.1080/23723556.2016.1164280.

70. Pacold ME, Brimacombe KR, Chan SH, Rohde JM, Lewis CA, Swier LJ, et al. A PHGDH inhibitor reveals coordination of serine synthesis and one-carbon unit fate. Nat Chem Biol 2016;12(6):452–8 doi 10.1038/nchembio.2070.

71. Weinstabl H, Treu M, Rinnenthal J, Zahn SK, Ettmayer P, Bader G, et al. Intracellular Trapping of the Selective Phosphoglycerate Dehydrogenase (PHGDH) Inhibitor BI-4924 Disrupts Serine Biosynthesis. J Med Chem 2019;62(17):7976–97 doi 10.1021/acs.jmedchem.9b00718.

72. Mullarky E, Xu J, Robin AD, Huggins DJ, Jennings A, Noguchi N, et al. Inhibition of 3-phosphoglycerate dehydrogenase (PHGDH) by indole amides abrogates de novo serine synthesis in cancer cells. Bioorg Med Chem Lett 2019;29(17):2503–10 doi 10.1016/j.bmcl.2019.07.011.

73. Riscoe MK, Brouns MC, Fitchen JH. Purine metabolism as a target for leukemia chemotherapy. Blood Rev 1989;3(3):162–73 doi 10.1016/0268-960x(89)90013-1.

74. Yamauchi T, Miyawaki K, Semba Y, Nakao F, Sugio T, Sasaki K, et al. Genome-Wide CRISPR Screen Identifies PAICS, An Enzyme Involved in De Novo Purine Synthesis, As a Potential Target for AML Therapy. Clinical Lymphoma, Myeloma and Leukemia 2019;19:S216–S7 doi 10.1016/j.clml.2019.07.088.

