## Supplementary Figures and Materials and Methods for "Reprogramming of serine metabolism is an actionable vulnerability in *FLT3*-*ITD* driven acute myeloid leukaemia"

### Supplementary Materials

This file includes the following supplementary materials:

- Supplementary Figures 1-6, with accompanying Figure legends;
- Supplementary Materials and Methods.

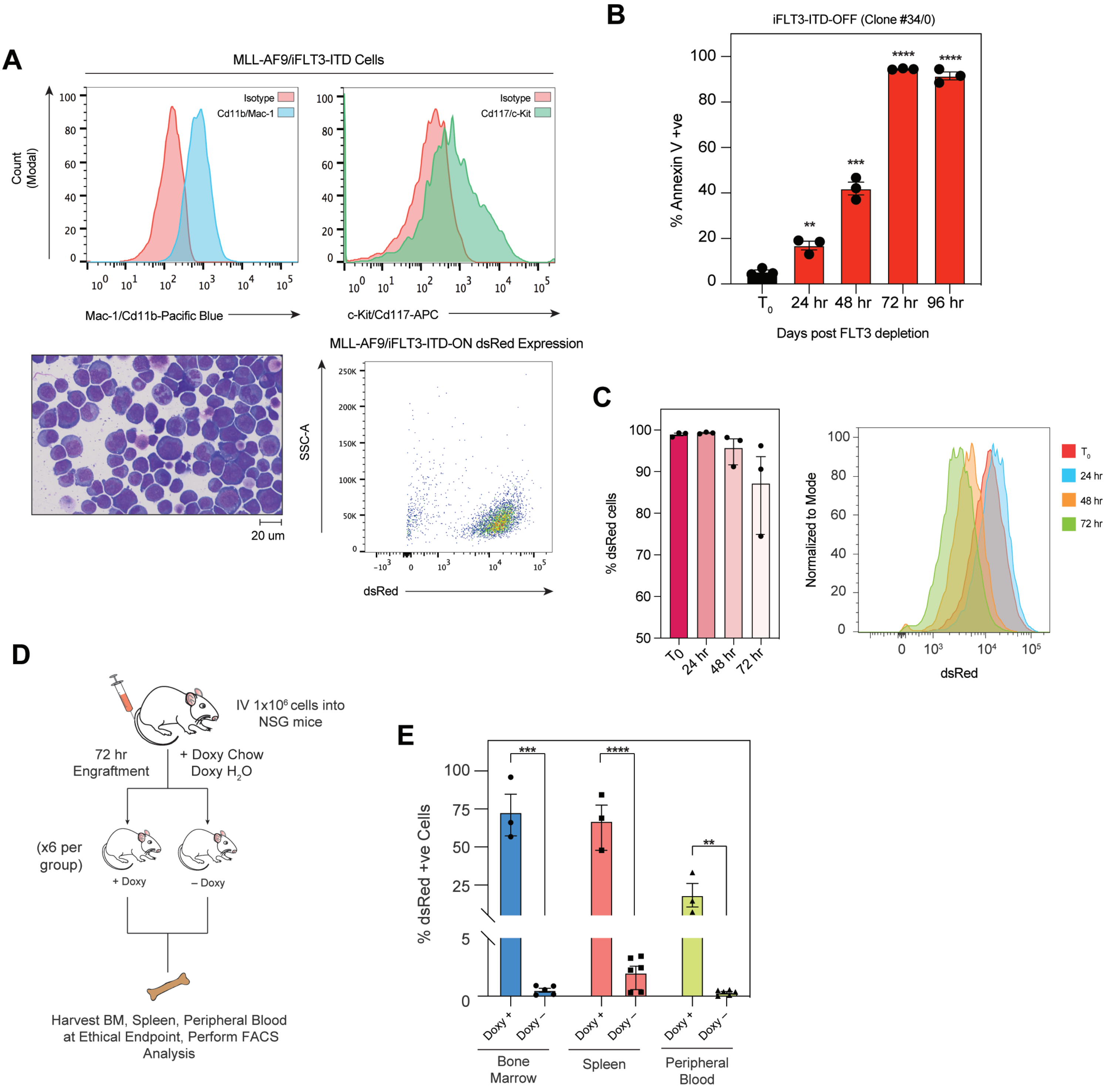


Supplementary Figure 1

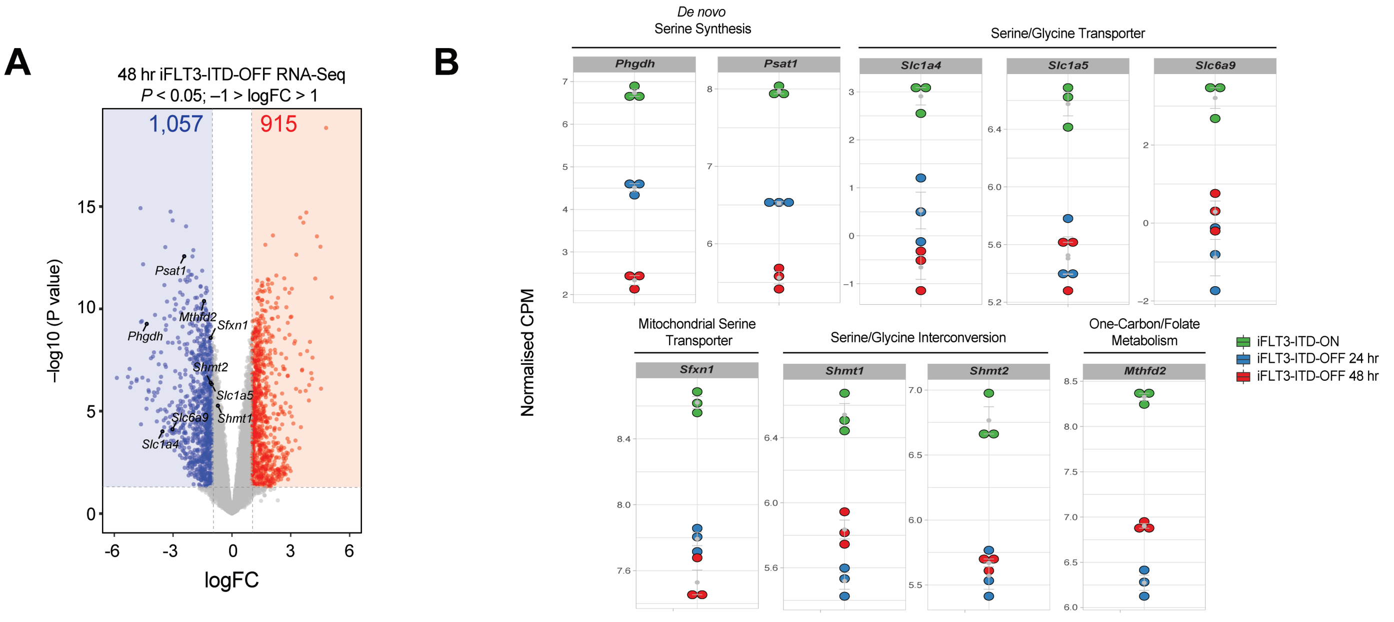


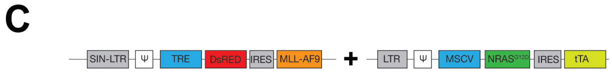


+ Doxycycline (24hr)

| Gene | logFC | Adj. *p*-value | Significant?# |
|---|---|---|---|
| *Phgdh* | 0.214 | 0.036 | Yes |
| *Psat1* | 0.288 | 0.0005 | Yes |
| *Slc1a4* | -0.327 | 0.443 | No |
| *Slc1a5* | 0.292 | 0.015 | Yes |
| *Slc6a9* | NE | NE | No |
| *Sfxn1* | -0.181 | 0.025 | Yes |
| *Shmt1* | -0.085 | 0.591 | No |
| *Shmt2* | 0.134 | 0.370 | No |
| *Mthfd2* | 0.179 | 0.052 | No |

*NE = not expressed/detected
#Assuming significant = $p < 0.05$

Supplementary Figure 2

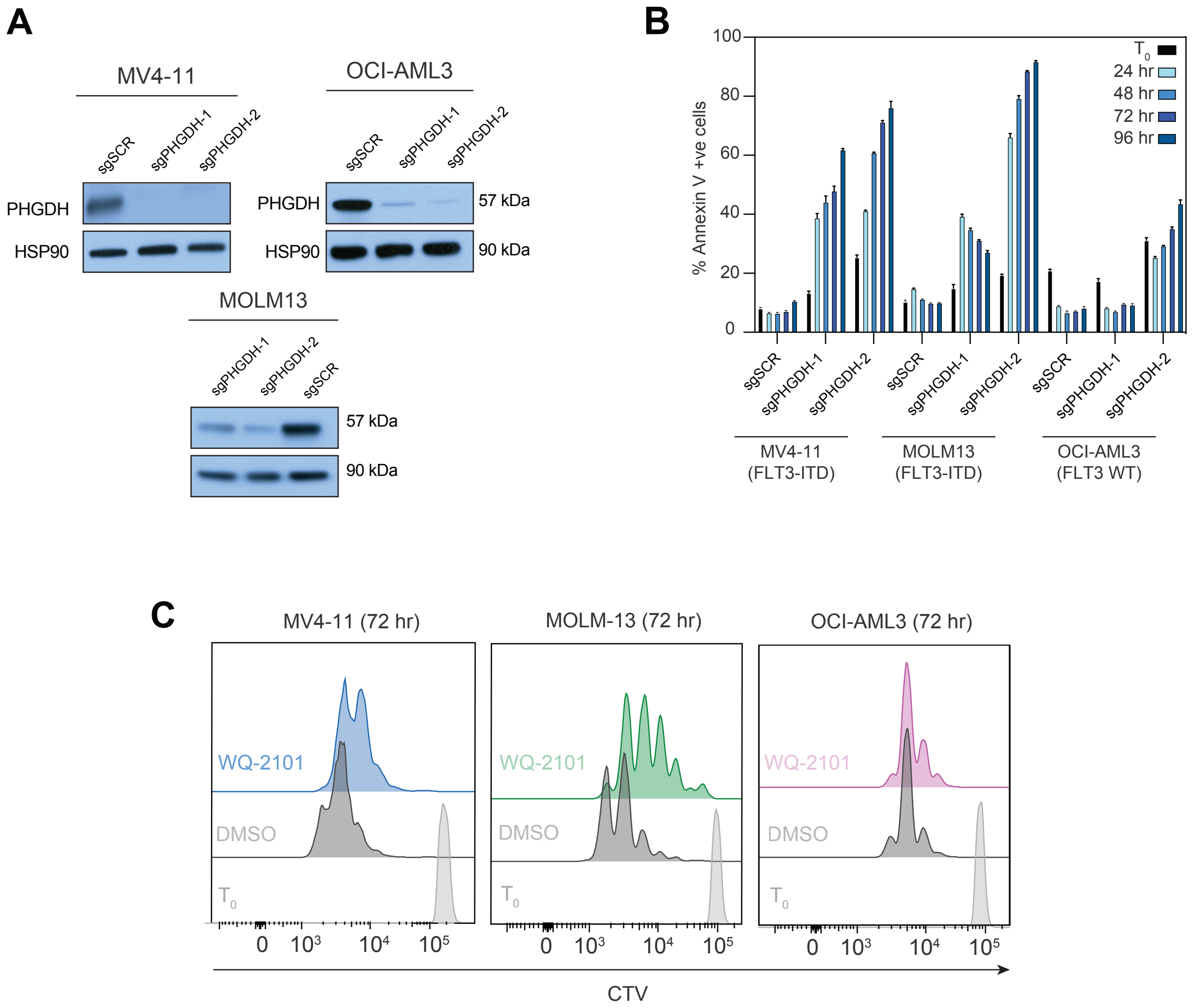


Supplementary Figure 3

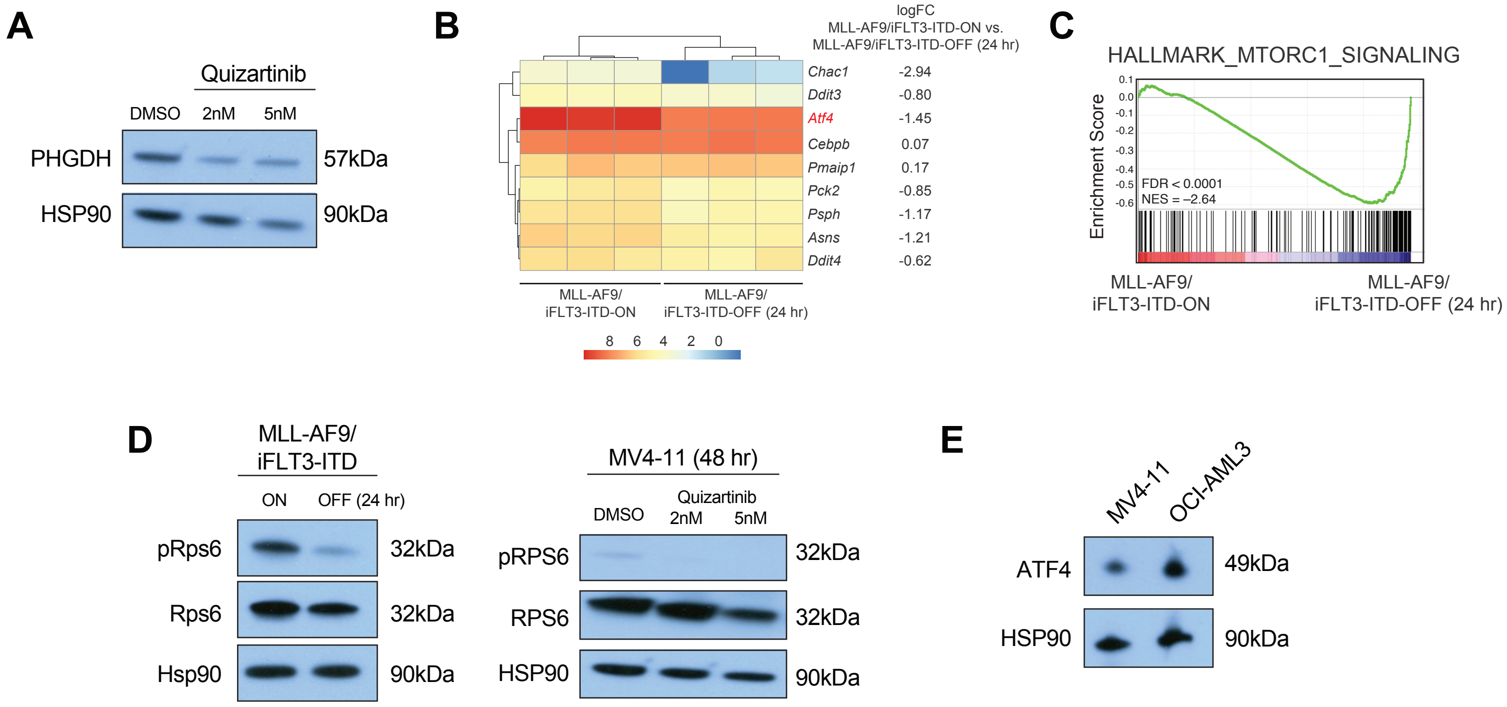


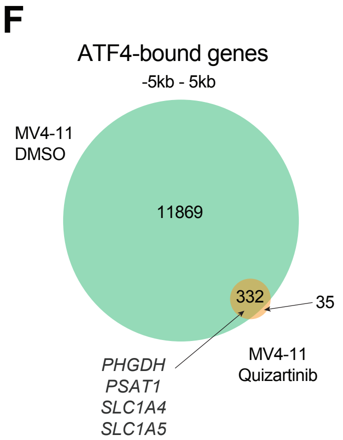


**G**

*De novo* motif –
ATF4 ChIP-Seq; MV4-11 DMSO

| Transcription Factor | Motif | Log p-value | % Targets | % Back-ground | Rank |
|---|---|---|---|---|---|
| Atf4 | ATTGCATCAC | -1.34e5 | 50.54 | 1.75 | 1 |
| Nr2f6 | GTCAAAGGAGAT | -3.75e3 | 1.66 | 0.07 | 3 |

*De novo* motif –
ATF4 ChIP-Seq; MV4-11 Quizartinib

| Transcription Factor | Motif | Log p-value | % Targets | % Back-ground | Rank |
|---|---|---|---|---|---|
| Atf4 | TGATGCAATG | -3.57e3 | 43.32 | 0.58 | 1 |
| CEBP:AP1 | AGAGTGTTTCAA | -7.03e2 | 7.88 | 0.07 | 4 |

Supplementary Figure 4

# A

#### Non-Essential Amino Acids

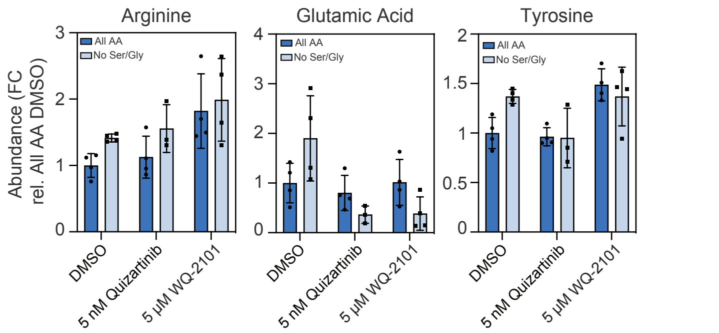


| Condition 1 | Condition 2 | Significance |
|---|---|---|
| DMSO AA | Quizartinib AA | ns |
| DMSO AA | WQ-2101 AA | p = 0.0317 |
| DMSO No Ser | Quizartinib No Ser | ns |
| DMSO No Ser | WQ-2101 No Ser | ns |
| DMSO AA | DMSO No Ser | p = 0.0047 |
| Quizartinib AA | Quizartinib No Ser | ns |
| WQ-2101 AA | WQ-2101 No Ser | ns |

| Condition 1 | Condition 2 | Significance |
|---|---|---|
| DMSO AA | Quizartinib AA | ns |
| DMSO AA | WQ-2101 AA | ns |
| DMSO No Ser | Quizartinib No Ser | p = 0.0309 |
| DMSO No Ser | WQ-2101 No Ser | p = 0.0168 |
| DMSO AA | DMSO No Ser | ns |
| Quizartinib AA | Quizartinib No Ser | ns |
| WQ-2101 AA | WQ-2101 No Ser | ns |

| Condition 1 | Condition 2 | Significance |
|---|---|---|
| DMSO AA | Quizartinib AA | ns |
| DMSO AA | WQ-2101 AA | p = 0.0051 |
| DMSO No Ser | Quizartinib No Ser | p = 0.0391 |
| DMSO No Ser | WQ-2101 No Ser | ns |
| DMSO AA | DMSO No Ser | p = 0.0052 |
| Quizartinib AA | Quizartinib No Ser | ns |
| WQ-2101 AA | WQ-2101 No Ser | ns |

# B

#### Essential Amino Acids

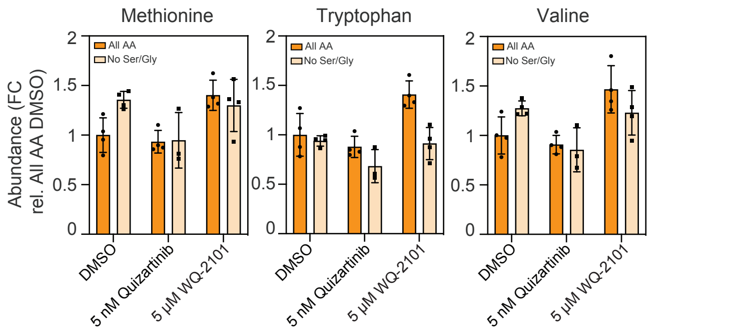


| Condition 1 | Condition 2 | Significance |
|---|---|---|
| DMSO AA | Quizartinib AA | ns |
| DMSO AA | WQ-2101 AA | p = 0.0131 |
| DMSO No Ser | Quizartinib No Ser | p = 0.0369 |
| DMSO No Ser | WQ-2101 No Ser | ns |
| DMSO AA | DMSO No Ser | p = 0.0103 |
| Quizartinib AA | Quizartinib No Ser | ns |
| WQ-2101 AA | WQ-2101 No Ser | ns |

| Condition 1 | Condition 2 | Significance |
|---|---|---|
| DMSO AA | Quizartinib AA | ns |
| DMSO AA | WQ-2101 AA | p = 0.0192 |
| DMSO No Ser | Quizartinib No Ser | p = 0.0317 |
| DMSO No Ser | WQ-2101 No Ser | ns |
| DMSO AA | DMSO No Ser | ns |
| Quizartinib AA | Quizartinib No Ser | ns |
| WQ-2101 AA | WQ-2101 No Ser | p = 0.0035 |

| Condition 1 | Condition 2 | Significance |
|---|---|---|
| DMSO AA | Quizartinib AA | ns |
| DMSO AA | WQ-2101 AA | p = 0.0220 |
| DMSO No Ser | Quizartinib No Ser | p = 0.0155 |
| DMSO No Ser | WQ-2101 No Ser | ns |
| DMSO AA | DMSO No Ser | p = 0.0352 |
| Quizartinib AA | Quizartinib No Ser | ns |
| WQ-2101 AA | WQ-2101 No Ser | ns |

# C

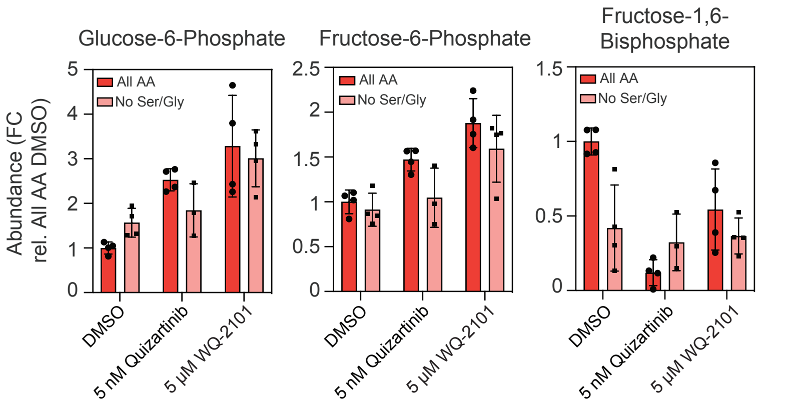


| Condition 1 | Condition 2 | Significance |
|---|---|---|
| DMSO AA | Quizartinib AA | p < 0.0001 |
| DMSO AA | WQ-2101 AA | p = 0.0073 |
| DMSO No Ser | Quizartinib No Ser | ns |
| DMSO No Ser | WQ-2101 No Ser | p = 0.0068 |
| DMSO AA | DMSO No Ser | p = 0.0172 |
| Quizartinib AA | Quizartinib No Ser | ns |
| WQ-2101 AA | WQ-2101 No Ser | ns |

| Condition 1 | Condition 2 | Significance |
|---|---|---|
| DMSO AA | Quizartinib AA | p = 0.0021 |
| DMSO AA | WQ-2101 AA | p = 0.0011 |
| DMSO No Ser | Quizartinib No Ser | ns |
| DMSO No Ser | WQ-2101 No Ser | p = 0.0170 |
| DMSO AA | DMSO No Ser | ns |
| Quizartinib AA | Quizartinib No Ser | ns |
| WQ-2101 AA | WQ-2101 No Ser | ns |

| Condition 1 | Condition 2 | Significance |
|---|---|---|
| DMSO AA | Quizartinib AA | p < 0.0001 |
| DMSO AA | WQ-2101 AA | p = 0.0190 |
| DMSO No Ser | Quizartinib No Ser | ns |
| DMSO No Ser | WQ-2101 No Ser | ns |
| DMSO AA | DMSO No Ser | p = 0.0086 |
| Quizartinib AA | Quizartinib No Ser | ns |
| WQ-2101 AA | WQ-2101 No Ser | ns |

# D

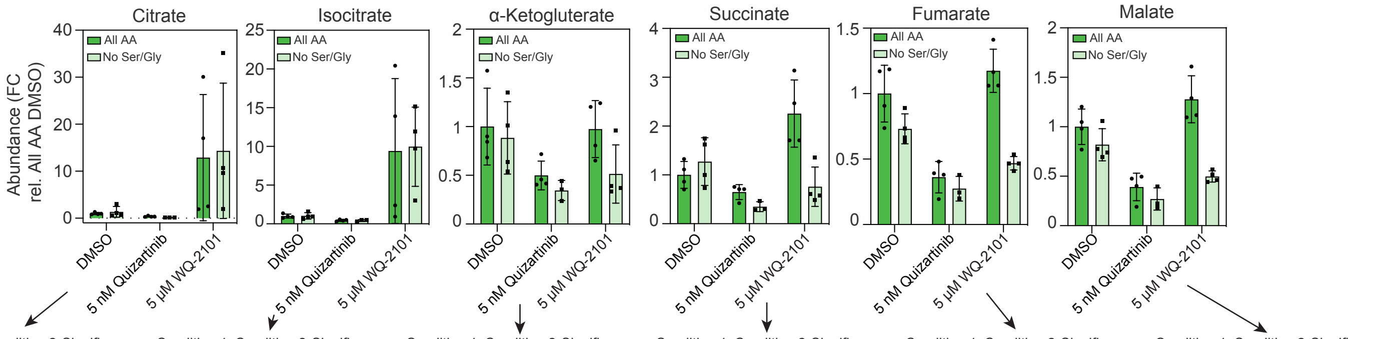


| Condition 1 | Condition 2 | Significance |
|---|---|---|
| DMSO AA | Quizartinib AA | p = 0.0039 |
| DMSO AA | WQ-2101 AA | ns |
| DMSO No Ser | Quizartinib No Ser | ns |
| DMSO No Ser | WQ-2101 No Ser | ns |
| DMSO AA | DMSO No Ser | ns |
| Quizartinib AA | Quizartinib No Ser | p = 0.0351 |
| WQ-2101 AA | WQ-2101 No Ser | ns |

| Condition 1 | Condition 2 | Significance |
|---|---|---|
| DMSO AA | Quizartinib AA | p = 0.0085 |
| DMSO AA | WQ-2101 AA | ns |
| DMSO No Ser | Quizartinib No Ser | ns |
| DMSO No Ser | WQ-2101 No Ser | p = 0.0132 |
| DMSO AA | DMSO No Ser | ns |
| Quizartinib AA | Quizartinib No Ser | ns |
| WQ-2101 AA | WQ-2101 No Ser | ns |

| Condition 1 | Condition 2 | Significance |
|---|---|---|
| DMSO AA | Quizartinib AA | ns |
| DMSO AA | WQ-2101 AA | ns |
| DMSO No Ser | Quizartinib No Ser | ns |
| DMSO No Ser | WQ-2101 No Ser | ns |
| DMSO AA | DMSO No Ser | ns |
| Quizartinib AA | Quizartinib No Ser | ns |
| WQ-2101 AA | WQ-2101 No Ser | ns |

| Condition 1 | Condition 2 | Significance |
|---|---|---|
| DMSO AA | Quizartinib AA | ns |
| DMSO AA | WQ-2101 AA | p = 0.0146 |
| DMSO No Ser | Quizartinib No Ser | p = 0.0253 |
| DMSO No Ser | WQ-2101 No Ser | ns |
| DMSO AA | DMSO No Ser | ns |
| Quizartinib AA | Quizartinib No Ser | p = 0.0340 |
| WQ-2101 AA | WQ-2101 No Ser | p = 0.0094 |

| Condition 1 | Condition 2 | Significance |
|---|---|---|
| DMSO AA | Quizartinib AA | p = 0.0021 |
| DMSO AA | WQ-2101 AA | ns |
| DMSO No Ser | Quizartinib No Ser | p = 0.0025 |
| DMSO No Ser | WQ-2101 No Ser | p = 0.0057 |
| DMSO AA | DMSO No Ser | ns |
| Quizartinib AA | Quizartinib No Ser | ns |
| WQ-2101 AA | WQ-2101 No Ser | p = 0.0002 |

| Condition 1 | Condition 2 | Significance |
|---|---|---|
| DMSO AA | Quizartinib AA | p = 0.0017 |
| DMSO AA | WQ-2101 AA | ns |
| DMSO No Ser | Quizartinib No Ser | p = 0.0042 |
| DMSO No Ser | WQ-2101 No Ser | p = 0.0097 |
| DMSO AA | DMSO No Ser | ns |
| Quizartinib AA | Quizartinib No Ser | ns |
| WQ-2101 AA | WQ-2101 No Ser | p = 0.0007 |

Supplementary Figure 5

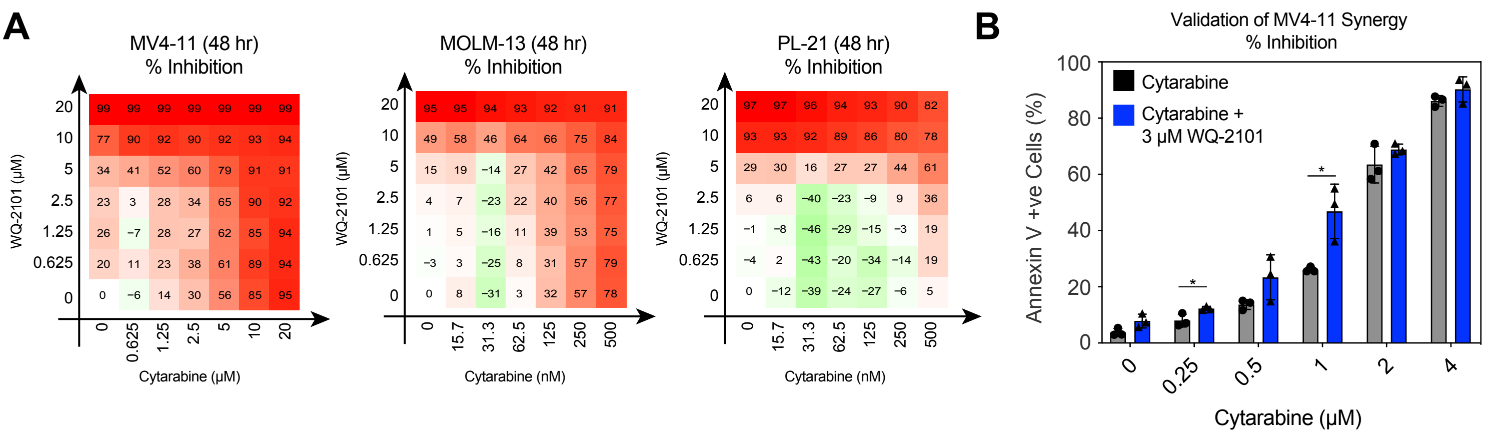


**C**

| Primary AML Patient Clinical Information | | |
|---|---|---|
| **Sample ID#** | AML-01-226-2014 | AML-01-309-2014 |
| **Sex; Age at time of marrow** | Male; 73 years | Male; 66 years |
| ***De novo* or therapy AML** | *De novo* | *De novo* |
| **AML type** | AML without maturation, intermediate prognosis karyotype | AML with myelodysplasia-related changes |
| **Sample type** | 5+2 chemotherapy refractory | Treatment naïve |
| **Mutational profile** | No *NPM1* or other mutations detected on targeted panel | *DNMT3A* R882S; no *NPM1* or other mutations on a targeted panel |
| **Karyotype** | Unbalanced 11;19 translocation<br><br>FISH did not identify any KMT2A (11q23) signal rearrangements in 200 cells scored, however three KMT2A signals were seen in 117/200 (58.5%) cells scored, confirming 11q translocation on 19p. | Normal |
| **FLT3-ITD to WT allelic ratio** | 1.16; mutational load 53.6% | 0.36; mutational load 26.6% |

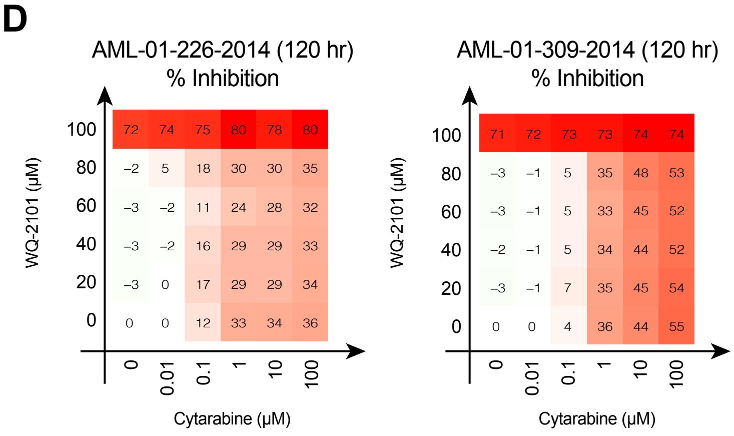


Supplementary Figure 6

#### SUPPLEMENTARY FIGURES

**Supplementary Figure 1. A genetically engineered mouse model of MLL-rearranged, FLT3-ITD AML reveals FLT3-ITD is essential for leukaemia survival.**

**A.** Characterisation of primary MLL-AF9/iFLT3-ITD AML cells. Flow cytometry-based analysis demonstrated blasts expressed Mac-1/Cd11b and c-Kit/Cd117, consistent with MLL-rearranged AML as previously characterised (1). Isotype controls used in each experiment are denoted with red shading. May-Grünwald-Giemsa staining demonstrating histology consistent with AML blast cells. Scale is indicative of 20 μm. MLL-AF9/iFLT3-ITD cells can be monitored by dsRed expression via flow cytometry. **B.** A second independent primary MLL-AF9/iFLT3-ITD clone was harvested from mouse #34/0 and blasts were cultured *in vitro* in 1 μg/mL doxycycline. Cells were then washed and cultured in doxycycline-free medium to deplete FLT3-ITD expression for 96 hours, and percent apoptotic cells subsequently assessed via Annexin V flow cytometry every 24 hours. ** denotes $P < 0.01$, *** $P < 0.001$, **** $P < 0.0001$. Error bars are representative of ± s.d. and three independent biological replicates. **C.** dsRed expression post FLT3-ITD depletion was assessed at 24-hour intervals up to 72 hours via flow cytometry. Error bars are representative of ± s.d. and three independent biological replicates. **D.** Schematic representation of *in vivo* iFLT3-ITD depletion. Mice were IV injected with $1 \times 10^6$ MLL-AF9/iFLT3-ITD cells and placed on doxycycline containing chow and water. 72 hours post injection, half of the cohort was placed on normal chow and water to deplete FLT3-ITD transgene expression ($n$ = 6 mice per group). **E.** At endpoint, MLL-AF9/iFLT3-ITD-dsRed-positive leukaemic blast burden was assessed in bone marrow, spleen and peripheral blood compartments via flow cytometry. ** denotes $P < 0.01$, *** $P < 0.001$, **** $P < 0.0001$. Error bars are representative of ± s.d. and 3-6 independent biological replicates.

**Supplementary Figure 2. *In vitro* and *in vivo* transcriptomics reveals *de novo* serine synthesis and serine uptake is regulated by FLT3-ITD.**

**A.** Volcano plot of differentially expressed genes 48 hours post FLT3-ITD depletion, highlighting genes involved in mediation of *de novo* serine synthesis, channel transporters, and one-carbon/folate metabolism in MLL-AF9/iFLT3-ITD cells. Suppressed differentially expressed genes (1,057) are denoted by blue, and enriched genes (915) by red shading. Significance was defined as $P < 0.05$ and $-1 > \log FC > 1$. **B.** Dot plots of normalised counts-per-million (CPM) gene expression values 24 and 48-hours post FLT3-ITD depletion of genes highlighted in Supplementary Figure 2A. Genes are grouped under biological/molecular function. Error bars denote ± s.d. and each condition was composed of 3 independent biological replicates. **C.** Table depicting logFC, adjusted *P*-value and significance of key genes suppressed upon MLL-AF9 depletion in a Tet-inducible model of MLL-AF9. Tet-inducible MLL-AF9/$NRAS^{G12D}$ cells were cultured in 1 μg/mL doxycycline to suppress expression of MLL-AF9 for 24 hours, and RNA extracted and 3'-RNA-sequenced. NE, not expressed/detected; significance $P < 0.05$.

**Supplementary Figure 3. Serine metabolism is a metabolic vulnerability in FLT3-ITD-driven AML.**

**A.** Efficiency of sgRNAs against PHGDH (sgPHGDH-1 or -2) or Scramble DNA (sgSCR) was validated via Western blotting in MV4-11, MOLM-13 and OCI-AML3 cells. HSP90 served as the loading control in all experiments. **B.** MV4-11, MOLM-13 and OCI-AML3 transduced with either two independent sgRNAs against PHGDH (sgPHGDH-1 or -2) or Scramble DNA (sgSCR) and were cultured in serine/glycine deprived medium for 96 hours, and cell viability determined via Annexin-V flow cytometry every 24 hours. Raw viability data of each timepoint is shown, and error bars are representative of ± s.d. and 3 independent biological replicates. **C.** MV4-11, MOLM-13 and OCI-AML3 cells were treated with DMSO, or 3 μM WQ-2101 and proliferation assessed via Cell Trace Violet 72 hours post dosing. Data is representative of 3 independent experiments.

**Supplementary Figure 4. A FLT3-ITD/mTORC1/ATF4 axis transcriptionally modulates serine synthesis and serine transporters in FLT3-mutant AML.**

**A.** MV4-11 cells were treated with DMSO, or two doses of quizartinib (2 nM or 5 nM) for 48 hours, and PHGDH expression assessed via Western blot. HSP90 served as the loading control. **B.** Heatmap of *Atf4 target* gene suppression post 24-hour FLT3-ITD depletion in murine MLL-AF9/iFLT3-ITD cells. All gene expression changes are denoted as logFC. **C.** Gene set enrichment (GSEA) plots depicting suppressed mTORC1 signalling after 24-hour FLT3-ITD depletion in murine MLL-AF9/iFLT3-ITD cells, FDR < 0.0001, NES = -2.64. **D.** Murine MLL-AF9/iFLT3-ITD cells were cultured in the absence of doxycycline for 24 hours to deplete FLT3-ITD, and phospho- and total Rps6 assessed via Western blot. The experiment was repeated in MV4-11 cells treated with either DMSO, or two doses of quizartinib (2 nM or 5 nM) for 48 hours. HSP90 served as the loading control in both experiments. **E.** Baseline ATF4 protein levels were assessed in MV4-11 and OCI-AML3 cells via Western blot. HSP90 was the loading control. **F.** Venn diagram of ATF4-bound genes in MV4-11 DMSO and quizartinib treated cells conditions (-5kb to 5kb). **G.** *De novo* motif discovery on ATF4 bound sites in DMSO and quizartinib-treated conditions in MV4-11 cells.

**Supplementary Figure 5. FLT3-ITD regulates global serine metabolism and drives purine nucleotide biosynthesis.**

**A.** Fold changes in non-essential amino acids arginine, glutamic acid and tyrosine abundances, as measured by LC/MS-MS, in DMSO-treated MV4-11 cells versus MV4-11 cells treated with 5 nM quizartinib or 5 μM WQ-2101 for 24 hours in all amino acid (AA)-containing RPMI medium, or serine/glycine-depleted medium. Tables containing statistical analyses for each condition are below each graph, and calculated via Student's *t*-test. Error bars are indicative of ± s.d. and 4 biological replicates. **B.** Fold changes in essential amino acids methionine, tryptophan and valine abundances, as measured by LC/MS-MS, in DMSO-treated MV4-11 cells versus MV4-11 cells treated with 5 nM quizartinib or 5 μM WQ-2101 for 24 hours in all amino acid (AA)-containing RPMI medium, or serine/glycine-depleted medium. **C.** Fold changes in glucose-6-phosphate, fructose-6-phosphate and fructose-1,6-bisphosphate abundances, as measured by LC/MS-MS, in DMSO-treated MV4-11 cells versus MV4-11 cells treated with 5 nM quizartinib or 5 μM WQ-2101 for 24 hours in all amino acid (AA)-containing RPMI medium, or serine/glycine-depleted medium. Tables containing statistical analyses for each condition are below each graph, and calculated via Student's *t*-test. Error bars are indicative of ± s.d. and 4 biological replicates. **D.** Fold changes in citrate, isocitrate, α-ketoglutarate, succinate, fumarate and malate abundances, as measured by LC/MS-MS, in DMSO-treated MV4-11 cells versus MV4-11 cells treated with 5 nM quizartinib or 5 μM WQ-2101 for 24 hours in all amino acid (AA)-containing RPMI medium, or serine/glycine-depleted medium. Tables containing statistical analyses for each condition are below each graph, and calculated via Student's *t*-test. Error bars are indicative of ± s.d. and 4 biological replicates.

**Supplementary Figure 6. Inhibition of *de novo* serine synthesis sensitizes FLT3-ITD-driven AMLs to cytarabine.**

**A.** MV4-11, MOLM-13 and PL-21 cells were treated with escalating concentrations of WQ-2101 and cytarabine for 48 hours to determine the effect on viability as determined by Cell Titer Glo. The mean of three biological replicates was used to determine each data point, and was normalized to DMSO controls. **B.** MV4-11 cells were treated with escalating doses of cytarabine and/or 3 μM WQ-2101 for 48 hours, and cell viability assessed using a non-ATP-based readout (Annexin V flow cytometry). Error bars are indicative of ± s.d. and 3 biological replicates. * denotes $P < 0.05$ as determined by Student's *t*-test. **C.** Clinical information summary of primary AML patient samples. **D.** Cells from two independent primary AMLs, AML-01-226-2014 and AML-01-309-2014, were treated with escalating concentrations of WQ-2101 and cytarabine for 120 hours to determine the effect on cell viability as determined by Sytox Blue flow cytometry. Data is indicative of one experiment.

#### SUPPLEMENTARY TABLES

**Supplementary Table S1**: Differentially expressed genes 24 hours post FLT3-ITD depletion in MLL-AF9/iFLT3-ITD-OFF cells.

**Supplementary Table S2**: Differentially expressed genes 48 hours post FLT3-ITD depletion in MLL-AF9/iFLT3-ITD-OFF cells.

**Supplementary Table S3:** Differentially expressed genes 24 hours post MLL-AF9 depletion in TRE-tight–DsRED–IRES–MLL-AF9/MSCV–NRAS–IRES–tTA cells.

**Supplementary Table S4:** Differentially expressed genes 48 hours post *in vivo* FLT3-ITD depletion in TET-FLT3-OFF cells.

**Supplementary Table S5:** Differentially expressed genes 24 hours post 5 nM quizartinib treatment in MV4-11 cells.

**Supplementary Table S6:** 84 gene overlap between MLL-AF9/iFLT3-ITD-OFF (24 hr) cells and 24 hr quizartinib treatment in MV4-11 cells.

**Supplementary Table S7:** 64 gene overlap of ATF4-bound target genes directly transcriptionally regulated by ATF4.

#### SUPPLEMENTARY REFERENCES

1. Somervaille TCP, Cleary ML. Identification and characterization of leukemia stem cells in murine MLL-AF9 acute myeloid leukemia. Cancer Cell **2006**;10(4):257-68 doi https://doi.org/10.1016/j.ccr.2006.08.020.

#### Supplementary Materials and Methods

*Retroviral generation of TET-FLT3-ITD/MLL-AF9 cells and foetal liver transductions*

For generation of murine TET-FLT3-ITD/MLL-AF9 cells, FLT3-ITD was cloned into a TRE-tight–DsRED–IRES–Empty vector and MLL-AF9 was cloned into an MSCV–Luc2–IRES–Empty vector using standard cloning techniques, and as previously described by our laboratory (1). All transfections were performed in HEK-293T cells using polyethyleneimine (PEI) reagent (Sigma-Aldrich), and the pCL-Ampho retroviral packaging vector. Viral supernatant was collected at 48- and 72-hours post transfection, and spun onto Retronectin (Takhara)-coated tissue culture plates at 2000*g* for 1 hour in either in MLL-AF9 alone or 1:1 MLL-AF9/FLT3-ITD virus conditions. 1 x $10^6$ 13.5 embryonic mouse day foetal liver cells were harvested from mice endogenously expressing the CAG-rtTA3 transgene, and subsequently spun onto viral-coated plates at 500*g* for 4 minutes, and allowed to colocalise with virus for 48 hours. *Ptprca* mice were then sub-lethally irradiated with 6.5 Gy gamma irradiation, allowed to rest for 4 hours, and then IV injected via tail vein with transduced cells. Leukaemias were allowed to engraft, and monitoring of disease was performed via luciferase bioluminescence imaging as described. At endpoint, mice were ethically euthanised, and AML blasts extracted from bone marrow and spleen. Cells from two independent clones, #21/0 and #34/0, were isolated from bone marrow and cultured in Anne Kelso modified DMEM as described, in 1 μg/mL doxycycline to maintain expression of the FLT3-ITD transgene. In all genetic depletion experiments of FLT3-ITD, cells were cultured for a minimum of 24 hours in doxycycline to stabilise transgene expression, prior to 2 washes with PBS to remove doxycycline from medium. Cells were subsequently cultured for the specified amount of time and analysed as described.

*In Vivo analysis*

All *in vivo* experiments performed in this study were approved by the Peter MacCallum Cancer Centre (PMCC) Animal Ethics Committee under ethics approval numbers E555 and E627. *Ptprca,* NSG, and C57BL/6J mice endogenously expressing the CAG-rtTA3 transgene were bred in-house (PMCC). For primary *Ptprca* recipient transplants, 4-6-week-old females were sub-lethally irradiated with 6.5 Gy whole-body gamma-ray irradiation, using a Gammacell 1000 Cs-137 irradiator (AECL). 1 x $10^6$ retrovirally-transduced foetal liver cells were then IV tail-vein injected, and mice placed on doxycycline-containing chow (Specialty Foods) and water (0.2% doxycycline hyclate (Sigma-Aldrich), 2% sucrose) until ethical endpoint (hunched, ruffled, impaired movements). For secondary recipients, 4-6-week-old female NSG mice were IV-injected with MLL-AF9/FLT3 cells as described above. At endpoint, blast cells were harvested from femur and tibial/fibial bone. For *in vivo* bioluminescence imaging, mice were injected with 50 mg/kg D-luciferin and imaged in an IVIS bioluminescence imager (Perkin-Elmer) to assess disease burden. Analysis and normalization of *in vivo* luminescence was performed using Living Image v3.2 and v4.2 software.

*Retroviral/lentiviral transductions and plasmid cloning*

For generation of Tet-inducible sgRNAs, two independent sgRNA sequences were utilised from the Brunello library (2) against *PHGDH* – sgPHGDH-1: *5'-GATGACATCAGCGGTCACCT* and sgPHGDH-2: *5'-GACACACCTACCTGTCGTGG*. sgRNAs were then cloned into the FgH1t_UTG Tet-inducible sgRNA vector (Addgene ID # 70183). HEK-293T cells were transfected using polyethyleneimine (PEI) reagent (Sigma-Aldrich) and the pMDLg/pRRE, pRSV-Rev and pVSV.G packaging vectors, and viral supernatant harvested 48- and 72-hours post transfection. Spin infections of constitutively expressing Cas9 MV4-11, MOLM-13 and OCI-AML3 cells (FUCas9-Cherry vector; Addgene ID # 70182) with harvested virus were performed at 1500*g* for 30 minutes with 4 mg/mL Sequa-brene reagent (Sigma-Aldrich) at room temperature. For generation of inducible shRNAs, two independent shRNA sequences were utilised against *ATF4* – shATF4-1: *5'-GCCTAGGTCTCTTAGATGATTCTCGAGAATCATCTAAGAGACCTAGGC* and shATF4-2: *5'-CCACTCCAGATCATTCCTTTACTCGAGTAAAGGAATGATCTGGAGTGG*. shRNAs were then cloned into the Tet-pLKO-puro Tet-inducible shRNA vector (Addgene ID # 21915) and transduced into MV4-11 cells as described above. shRNA-expressing MV4-11 cells were selected by culturing in 1 μg/mL puromycin (Sigma-Aldrich) for 5 days prior to experimental use, and activated by the addition of 200 ng/mL doxycycline to deplete ATF4.

*Metabolic rescue experiments*

For serine rescue experiments in sgPHGDH cells, 2.5 x $10^5$ cells were seeded in 24-well plates, and escalating doses of L-serine (Sigma-Aldrich) was supplemented into serine/glycine-free RPMI at seeding. Cells were incubated for 48 hours prior to cell death analysis via Annexin V staining as described in the main text. For metabolite rescue experiments in response to FLT3-ITD inhibition with QUIZARTINIB, 2.5 x $10^5$ exponentially-growing MV4-11 cells were cultured in 24-well plates for 48-72-hours as described, and cultured in the presence of 3 nM of QUIZARTINIB. Metabolites were then supplemented into medium at time of seeding as follows: For nucleoside rescue experiment, 150 μM purine cocktail mix (Adenine, Guanine in sterile water, heated to 37ºC prior to use – Sigma-Aldrich); 150 μM pyrimidine cocktail mix (Cytosine, Thymine, Uridine in sterile water – Sigma-Aldrich); 150 μM EmbryoMax Nucleosides (Merck-Millipore, Cat # ES-008-D) and 286 μM L-serine (Sigma-Aldrich). For metabolite rescue experiments, 500 μM sodium pyruvate (Sigma-

Aldrich); 15 mM sodium L-lactate (Sigma-Aldrich); 150 μM hypoxanthine (6-hydroxypurine – Sigma-Aldrich); nucleoside and serine concentration and source as above. Cell viability was assessed via Annexin V flow cytometry as described above.

*Serine/glycine-free RPMI*

To generate serine/glycine free RPMI 1640 otherwise identical to the commercial medium utilised in all other experiments in this study (Gibco 11875), we synthesised medium as per manufacturer's protocol (https://www.thermofisher.com/au/en/home/technical-resources/media-formulation.114.html), and omitted the addition of serine and glycine in the final formulation. At the time of culture, we added 10% or 20% dialysed FBS prepared as previously described (3). All other medium conditions, such as antibiotic and Glutamax supplementation was identical to regular culture medium (discussed in *Cell lines and culture* section).

*Tritiated serine labelling*

MV4-11, MOLM-13, and OCI-AML3 cells were cultured in full-serine/glycine RPMI with DMSO, 5 nM QUIZARTINIB, or 5 μM WQ-2101 for 24 hours. $2 \times 10^6$ cells were harvested per condition and washed twice with PBS, prior to resuspension and labelling in serine-free RPMI supplemented with 250nM of L-[$^3$H(G)]-Serine (Perkin Elmer, Cat # NET248250UC) for 30 mins. Post labelling, cells were washed twice with PBS, and a whole-cell lysate was obtained by resuspending cells in Laemmli lysis buffer. After boiling the lysate for 5 minutes at 95ºC, an equivalent volume of tritiated lysate was resuspended in Optiphase HiSafe 3 liquid scintillation cocktail (Perkin Elmer), and beta decay emissions detected with a Tri-Carb 2910 TR liquid scintillation analyser (Perkin Elmer). Data was analysed using QuantaSmart v4.02 software (Perkin Elmer). All experiments were performed using three biological replicates, and background absorbance was subtracted from each sample as appropriate.

*Primary and secondary antibodies*

The following antibodies were utilised in this study. Manufacturer, catalogue number, dilution and application for each antibody are provided.

**Primary Antibodies**

| Antibody | Manufacturer | Catalogue # | Dilution | Application |
|---|---|---|---|---|
| Phospho-FLT3 (Tyr842) (10A8) Rabbit mAb | Cell Signaling | 4577S | 1:500 | WB |
| FLT3 (8F2) Rabbit mAb | Cell Signaling | 3462S | 1:500 | WB |

| | | | | |
|---|---|---|---|---|
| Phospho-STAT5 (Tyr694) Rabbit mAb | Cell Signaling | 9351S | 1:1000 | WB |
| STAT5 Rabbit mAb | Cell Signaling | 9363S | 1:1000 | WB |
| PARP Rabbit mAb | Cell Signaling | 9542L | 1:1000 | WB |
| Caspase-3 Mouse Ab | BD Biosciences | 611048 | 1:1000 | WB |
| PHGDH Rabbit Ab | Sigma-Aldrich | HPA021241 | 1:1000 | WB |
| PSAT1 Rabbit Ab | Novus Biologicals | NBP1-55368 | 1:1000 | WB |
| ATF4 (D4B8) Rabbit mAb | Cell Signaling | 11815S | 1:1000<br>13 μL/ChIP | WB<br>ChIP |
| RNA Pol II (CTD4H8) Mouse mAb | Merck Millipore | 05-623 | 6 μL/ChIP | ChIP |
| Phospho-S6 Ribosomal Protein (Ser240/244) Rabbit Ab | Cell Signaling | 2215S | 1:1000 | WB |
| S6 Ribosomal Protein (54D2) Mouse mAb | Cell Signaling | 2317S | 1:1000 | WB |
| Phospho-Histone H2A.X (Ser139) (20E3) Rabbit mAb | Cell Signaling | 9718S | 1:1000 | WB |
| β-Actin (AC-74) Mouse mAb | Sigma-Aldrich | A2228 | 1:5000 | WB |
| HSP90 (AC88) Mouse mAb | Enzo Life Sciences | ADI-SPA-830 | 1:3000 | WB |

**Secondary Antibodies**

| Antibody | Manufacturer | Catalogue # | Dilution | Application |
|---|---|---|---|---|
| Swine Anti-Rabbit Immunoglobulin HRP Ab | Dako | P0127 | 1:3000 | WB |
| Rabbit Anti-Mouse Immunoglobulin HRP Ab | Dako | P0260 | 1:3000 | WB |

*QuantSeq 3'-RNA sequencing and analysis*

Cells were cultured/harvested as described, and RNA extracted using TRIzol reagent (Thermo Fisher Scientific) and the Direct-Zol RNA Miniprep kit (Zymo Research) as per manufacturer's instructions. The QuantSeq 3'RNA-seq Library Prep Kit for Illumina (Lexogen) was then utilised from 500 ng of RNA to generate libraries as per manufacturer's instructions. Libraries were then sequenced on the Illumina NextSeq 500, with single-end 75bp reads to a depth of 15M reads per sample. Sequencing reads were subsequently trimmed at the 5'-end using CutAdapt (v1.14) software (4) to remove random primers introduced during library preparation. 3' ends were also trimmed to remove poly-A-tail derived reads. Reads were then mapped to reference genomes (mm10 for all mouse sequencing, hg19 for human) using HISAT2 (v2.1.0) software (5). Counting of reads was conducted using featureCounts, a component of the subread package (v1.5.2) (6). Differential gene expression was performed using the Voom-Limma methodology to determine statistical significance (7,8). All RNA-seq analysis figures were generated using R (v3.6.1) software. GO Term analysis was performed using ToppGene software using term sets as described (9). Gene Set Enrichment Analysis (GSEA) software (v3.0) was used for identification of enriched gene sets, using the MSigDB KEGG and Hallmarks datasets (10).

*Chromatin immunoprecipitation library preparation and sequencing analysis*
MV4-11 and OCI-AML3 cells (60 x $10^6$ per condition) were cultured in the presence or absence of 5 nM quizartinib for 24 hours, and pellets washed in ice-cold PBS. Cells were crosslinked with 1% formaldehyde for 10 minutes at room temperature and subsequently quenched with 1.25M glycine. Cells were washed three times with ice-cold PBS and lysed in ChIP lysis buffer (20 mM Tris-HCl, 150 mM $NaCl_2$, 2 mM EDTA, 1% IGEPAL CA-630 (Sigma-Aldrich) and 0.3% SDS in water) prior to sonication on a Covaris ultrasonicator at maximum power for 16 minutes to achieve an average DNA fragment size of 300-500bp. Immunoprecipitation reactions were performed in ChIP dilution buffer (20 mM Tris-HCl, 150 mM $NaCl_2$, 2 mM EDTA, 1% Triton-X, and phosphatase/protease inhibitors in water) overnight at 4ºC. A 1:1 ratio of protein A and G magnetic beads (Life Technologies) were used to bind crosslinked protein/DNA. Antibodies used in ChIP-seq experiments are outlined in the '*Primary and secondary antibodies*' section of this paper. Beads were then washed in increasingly high-salt concentration wash buffers and reverse crosslinked in the presence of Proteinase K (Zymo Research) prior to purification and concentration with the ChIP DNA Clean and Concentrator Kit (Zymo Research) per manufacturer's specifications.

Post elution and concentration of DNA, quantification was performed using the Qubit dsDNA HS Assay kit (Thermo Fisher), and ChIP-Seq libraries were prepared using the NEBNext Ultra II DNA Library Prep Kit for Illumina (New England BioLabs) as per manufacturer's instructions. Libraries were then size selected between 200bp and 500bp using a Pippin Prep 1.5% agarose cassette (Sage Science). Final library QC was performed on a Tape Station High Sensitivity DNA Analysis Kit (Agilent Technologies). For sequencing analysis of DNA immunoprecipitated with anti-ATF4 and anti-RNA Pol II antibodies, libraries were then pooled and sequenced on an Illumina NextSeq 500 and 20-25 million single-end 75bp reads were generated for each sample.

Raw sequencing reads were demultiplexed using bcl2fastq (v2.17.1.14) and FASTQ files generated and quality controlled with FASTQC (v0.11.5). Adaptor sequences were removed and reads aligned to the human reference genome (GRCh37/hg19) using Bowtie2 (v2.3.5). Samtools (v1.4.1) was used for processing of SAM and BAM files, from which Model-based Analysis of ChIP-Seq (MACS2, v2.2.5) peak calling was performed (11). TDF files were generated with IGVTools (v2.5.3) and ChIP-Seq tracks were visualised and figures generated using IGV software (v2.6.2). HOMER (v4.10) was used for quantification and annotation of the ChIP-Seq datasets, for determining peak scores of ATF4 target genes, and for *de novo* motif discovery on ATF4 bound sites. ATF4 genome-wide binding in MV4-11 and OCI-AML3 cells was visualised using the heatmap and profile functions within deeptools (v3.3.1) software.

*LC/MS-MS and metabolomics analysis*

MV4-11 cells were maintained in full growth medium and fresh medium (full or serine/glycine-free) was added at the time cells were treated with quizartinib or WQ-2101. Following a 24-hour treatment, 3 x $10^6$ MV4-11 cells were harvested by centrifugation, washed with normal saline and cell pellets were snap-frozen. For metabolite extraction, cell pellets were resuspended in 500 μL ice-cold MeOH:H2O (80:20) containing internal standards (13C-AMP, 13C-UMP, 13C Sorbitol, and 13C Valine) and vortexed. Samples were incubated on ice for 5 minutes, vortexed and debris was pelleted by centrifugation at 16,000 *g* for 10 minutes. The resulting supernatants were injected and analysed by hydrophilic interaction liquid chromatography (HILIC) and high-resolution mass spectrometry (Agilent 6545 LC/Q-TOF). Quality control checks were performed using QTOF MassHunter Quant software (Agilent) and metabolite peak calling was conducted using El-MAVEN analysis software (12). Data are presented as fold change in metabolite abundance relative to control.

*TCGA SingScore analysis*

To generate the serine synthesis pathway – one-carbon metabolism – purine signature (SSP-OC-Purine Signature), we utilised canonical genes associated with these pathways that were selected from our own transcriptional analyses in this paper, and supplemented from a study by Manning and colleagues (13). Importantly, we selected the most differentially-expressed purine biosynthesis genes from this study, while excluding those with minimal differential expression. A complete table of the genes utilised from each pathway are summarised in the table below. For interrogation of the LAML-TCGA dataset, we used the TCGAbiolinks R package (v2.15.3) to download and normalise the mRNA counts file using the edgeR method (8). FLT3 mutation status of each of the LAML samples was downloaded from cBioPortal. Samples were stratified based on FLT3 mutation vs FLT3 WT, and the singscore method was subsequently utilised to plot *FLT3* mRNA LogRPKM values against the signature score (14).

| **SSP** | **OC** | **Purine** |
|---|---|---|
| *PHGDH* | *DHFR* | *PRPS1* |
| *PSAT1* | *MTHFD1* | *PRPS2* |
| *PSPH* | *MTHFD1L* | *PPAT* |
| *SLC1A4* | *MTHFD2* | *GART* |
| *SLC1A5* | *SHMT1* | |
| *SLC6A9* | *SHMT2* | |
| | *GLDC* | |
| | *TYMS* | |

*Primary patient samples*

Primary patient samples used for *in vitro* synergy studies were obtained from the Alfred Hospital (Melbourne, Australia) after obtaining informed consent under Alfred Health Ethics Committee-approved guidelines. These studies were performed under Alfred Heath ethics approval number 42/20. Primary AML blasts were cultured in StemSpan SFEM (Stemcell Technologies) supplemented with 10 ng/mL IL-3, 10 ng/mL IL-6, 50 ng/mL SCF and 50 ng/mL FLT3L (all Peprotech) and 35 nM UM171 and 500 nM stemreginin (StemCell Technologies). No patient data other than age, sex and genomic profile were made available to non-clinical/authorised authors in this manuscript.

*In vitro synergy assays and analysis*

For analysis of *in vitro* synergy with MV4-11, MOLM-13 and PL-21 cells, 8,000 cells were seeded per well in a 384-well plate and incubated with escalating doses of WQ-2101 and cytarabine (AraC) in a checkerboard format for 48 hours with a D300e Digital Dispenser (Tecan Technologies). At endpoint, viability was assessed using Cell Titer Glo. Synergy was determined by converting raw absorbance values to % cell death for each condition and normalising to DMSO controls. Synergy was determined by utilising the SynergyFinder R package (v1.4.2) (15) using the Bliss synergy model. To ensure synergistic effects were not confounded by the ATP-based readout of Cell Titer Glo, a selection of combination therapy dose ranges was repeated *in vitro*, and Annexin V flow cytometry used as an alternative method for quantifying cell viability.

For primary patient synergy analyses, 50,000 AML cells were plated in each well of a 96-well plate, and cultured with escalating doses of WQ-2101 and cytarabine (AraC) to a maximum of 100 μM for 5 days in medium conditions as described above. At endpoint, cells were stained with 5 μM Sytox Blue (Invitrogen) and cell viability assessed via flow cytometry. Synergy analysis was performed as above.

*In Vivo Therapy Experiments*

For *in vivo* combination therapy experiments, 2.5 x $10^6$ MV4-11 cells transduced with a MSCV-Luc2-IRES-mCherry reporter were IV-injected into 6-8-week-old female NSG recipients. Therapy commenced after blast content of peripheral blood reached a threshold of ≥1% mCherry-positive cells. For *in vivo* combination assays, 5mg/kg WQ-2101 was i.p. injected into mice daily for 12 days, for a total of 13 doses of compound. 40mg/kg cytarabine (AraC) was i.p. injected on alternate days for a total of 6 doses of compound. After completion of therapy, mice were culled at ethical endpoint as described above. No censored events were recorded in this study, and all mice who received treatment were included in survival analysis.

#### SUPPLEMENTARY REFERENCES


1. Ghisi M, Kats L, Masson F, Li J, Kratina T, Vidacs E*, et al.* Id2 and E Proteins Orchestrate the Initiation and Maintenance of MLL-Rearranged Acute Myeloid Leukemia. Cancer Cell **2016**;30(1):59-74 doi 10.1016/j.ccell.2016.05.019.
2. Doench JG, Fusi N, Sullender M, Hegde M, Vaimberg EW, Donovan KF*, et al.* Optimized sgRNA design to maximize activity and minimize off-target effects of CRISPR-Cas9. Nat Biotechnol **2016**;34(2):184-91 doi 10.1038/nbt.3437.
3. Cantor JR, Abu-Remaileh M, Kanarek N, Freinkman E, Gao X, Louissaint A, Jr.*, et al.* Physiologic Medium Rewires Cellular Metabolism and Reveals Uric Acid as an Endogenous Inhibitor of UMP Synthase. Cell **2017**;169(2):258-72.e17 doi 10.1016/j.cell.2017.03.023.
4. Martin M. Cutadapt removes adapter sequences from high-throughput sequencing reads. 2011 **2011**;17(1):3 doi 10.14806/ej.17.1.200.
5. Kim D, Paggi JM, Park C, Bennett C, Salzberg SL. Graph-based genome alignment and genotyping with HISAT2 and HISAT-genotype. Nat Biotechnol **2019**;37(8):907-15 doi 10.1038/s41587-019-0201-4.
6. Liao Y, Smyth GK, Shi W. featureCounts: an efficient general purpose program for assigning sequence reads to genomic features. Bioinformatics **2014**;30(7):923-30 doi 10.1093/bioinformatics/btt656.
7. Law CW, Chen Y, Shi W, Smyth GK. voom: Precision weights unlock linear model analysis tools for RNA-seq read counts. Genome Biol **2014**;15(2):R29 doi 10.1186/gb-2014-15-2-r29.
8. Law CW, Alhamdoosh M, Su S, Dong X, Tian L, Smyth GK*, et al.* RNA-seq analysis is easy as 1-2-3 with limma, Glimma and edgeR. F1000Res **2016**;5 doi 10.12688/f1000research.9005.3.
9. Chen J, Bardes EE, Aronow BJ, Jegga AG. ToppGene Suite for gene list enrichment analysis and candidate gene prioritization. Nucleic Acids Res **2009**;37(Web Server issue):W305-11 doi 10.1093/nar/gkp427.
10. Liberzon A, Birger C, Thorvaldsdóttir H, Ghandi M, Mesirov JP, Tamayo P. The Molecular Signatures Database (MSigDB) hallmark gene set collection. Cell Syst **2015**;1(6):417-25 doi 10.1016/j.cels.2015.12.004.
11. Zhang Y, Liu T, Meyer CA, Eeckhoute J, Johnson DS, Bernstein BE*, et al.* Model-based analysis of ChIP-Seq (MACS). Genome Biol **2008**;9(9):R137 doi 10.1186/gb-2008-9-9-r137.
12. Agrawal S, Kumar S, Sehgal R, George S, Gupta R, Poddar S*, et al.* El-MAVEN: A Fast, Robust, and User-Friendly Mass Spectrometry Data Processing Engine for Metabolomics. Methods Mol Biol **2019**;1978:301-21 doi 10.1007/978-1-4939-9236-2_19.
13. Ben-Sahra I, Hoxhaj G, Ricoult SJH, Asara JM, Manning BD. mTORC1 induces purine synthesis through control of the mitochondrial tetrahydrofolate cycle. Science **2016**;351(6274):728-33 doi 10.1126/science.aad0489.
14. Foroutan M, Bhuva DD, Lyu R, Horan K, Cursons J, Davis MJ. Single sample scoring of molecular phenotypes. BMC Bioinformatics **2018**;19(1):404 doi 10.1186/s12859-018-2435-4.
15. Ianevski A, He L, Aittokallio T, Tang J. SynergyFinder: a web application for analyzing drug combination dose-response matrix data. Bioinformatics **2017**;33(15):2413-5 doi 10.1093/bioinformatics/btx162.
